# Stakeholder perspectives on the effectiveness of the Ifakara-Lupiro-Mang’ula Wildlife Management Area in Southern Tanzania

**DOI:** 10.1101/2024.11.12.623190

**Authors:** Lucia J. Tarimo, Deogratius R. Kavishe, Fidelma Butler, Gerry F. Killeen, Felister Mombo

**Affiliations:** Department of Forest and Environmental Economics, Sokoine University of Agriculture, P.O.Box 3011, Morogoro, Tanzania; Department of Environmental Health and Ecological Sciences, Ifakara Health Institute, Ifakara Branch, Off Mlabani Passage, P.O. Box 53 Ifakara, Morogoro, Tanzania; School of Biological Earth & Environmental Sciences, University College Cork, Cork, Republic of Ireland; Environmental Research Institute, University College Cork, Cork, Environmental Research Institute, Republic of Ireland

**Keywords:** conservation, wildlife, encroachments, local communities, livelihoods, community development, CBNRM, WMAs

## Abstract

In Tanzania, Wildlife Management Areas (WMAs) are established on village lands, usually adjacent to centrally managed core protected areas like national parks and game reserves, and managed in a devolved manner by local village authorities. WMAs are intended to conserve wildlife outside such core protected areas, while also providing opportunities for local communities to derive tangible benefits from wildlife and other natural resources. This study evaluates the perceived effectiveness of the Ifakara-Lupiro-Mang’ula (ILUMA) WMA in southern Tanzania among various stakeholders, focusing on its conservation, livelihoods and community development functions. Based on thematic analysis of perspectives shared by stakeholders at national, regional, district and village levels through in-depth interviews, focus group discussions and a public stakeholders meeting, the WMA was considered to have succeeded to only a very modest extent in achieving its intended goals. Essentially all participants narrated that the area is heavily encroached by human activities, including livestock grazing, agriculture, illegal fishing, meat poaching, deforestation, charcoal burning, timber harvesting and even permanent settlements. Contributing factors include a recently growing influx of agro-pastoralist immigrants, top-down political interference, financial constraints, financial mismanagement, limited resources for operations, lack of conservation education, investors or government support, and recent displacement of encroachment pressure from newly upgraded, centrally-managed protected areas nearby. To ensure future success and sustainability of the WMA, participants recommended enhancing stakeholder involvement and community participation in WMA management, improved collaboration with nearby centrally-managed protected areas for implementing operations, overhaul of the WMA constitution to reflect current best practices, building capacity among relevant village leaders and elected WMA representatives and initiating conservation education initiatives for the local community. Overall, the WMA should explore alternative income sources beyond tourism, to ensure direct benefits for the community member through sustainable, carefully-regulated access to natural resources, and resolve ongoing conflict over land use between the long-established villages that govern ILUMA and agro-pastoralists immigrants who have moved into the area more recently.

## 1. Introduction

In order to ensure sustainable conservation and equitable benefit from natural resources, there has been a shift to community based conservation (CBC) approaches in several African countries over the past three decades (Gibson & Marks, 1995; Songorwa, 1999) and, more broadly, community based natural resource management (CBNRM) (Fortmann et al., 2001; Nelson & Agrawal, 2008; Roe, 2011). Such decentralized strategies emerged after the failure of conventional “fortress conservation” models (Brockington, 2002), which excluded local community and largely denied them meaningful from benefits derived from such protected areas (Goldman, 2003; Songorwa et al., 2000). These strictly exclusionary approaches led to increased conflict between local communities and conservation authorities, ultimately compromising effectiveness of biodiversity protection (Lane, 2001). CBNRM is based on the premise that sustainable management of natural resources is most readily and fairly achievable when local communities have the authority to manage and benefit from the relevant resources (Kull, 2002; Murphree, 2004) and, with their invaluable traditional knowledge, are better equipped to conserve natural landscapes (Agrawal & Gibson, 1999). By integrating local communities as key stakeholders, CBNRM aims to achieve win-win outcome that promotes both sustainable conservation of natural resources and rural development by fostering effective decentralization of governance and management (Treue & Nathan, 2007).

In Africa, CBNRM models were initially developed in Southern and Eastern African countries, with some impressive early successes. Notable examples include Zimbambwe’s Communal Areas Management Programme for Indigenous Resources (CAMPFIRE), which emerged in the 1970s and 1980s as the first African CBNRM initiative regarding wildlife management (Fortmann et al., 2001; Martin, 1986), the Namibian Conservancy Model, and Zambia Administrative Management Design (ADMADE), (Balint, 2006; USAID, 2013). Despite their imperfections and limitations (Chisanga, 2016; Dressler et al., 2010; Garner, 2012; Hulme & Murphree, 2001; Jones, 2010), these initiatives have nevertheless demonstrated success in achieving natural resource conservation goals, while also improving local livelihoods (Mbaiwa, 2004; Shackelton & Campbell, 2000).

In Tanzania, the CBNRM approach has been applied through several devolved institutional models, including Community Based Forest Management (CBFM), pioneered by establishment of Duru Haytemba community forest (Kajembe et al., 2006). Wildlife Management Areas (WMAs) represent another model drawing lessons from this pioneering initiative, where villages set aside land for wildlife conservation under their own local management (United Republic of Tanzania for Ministry of Natural Resources and Tourism, 1998). WMAs are intended to generate revenue for local communities through tourism and other complementary income generation activities (Baldus & Cauldwell, 2004; Kiss, 2004), while also preserving extensive and interconnected habitats for wildlife protection (United Republic of Tanzania for Ministry of Natural Resources and Tourism, 1998). Indeed, this rationale for the formation of WMAs emerged during the 1990s (USAID, 2013), driven by the need to protect areas large enough to facilitate extensive migration routes and populations of sparsely distributed animals that are big enough to have adequate genetic diversity. In 1998, the Tanzanian government formulated a Wildlife Policy (revised in 2007) that established the legal framework for WMA establishment, aiming to promote CBNRM through decentralized local management and ownership of wildlife resources (United Republic of Tanzania for Ministry of Natural Resources and Tourism, 1998). The primary goal of the WMA institutional model is to engage rural communities in collectively managing wildlife and other natural resources within village lands on a sustainable basis, thereby conserving important wildlife areas such as migration corridors, buffer zones for adjacent parks and reserves, and important habitat used by animals in particular seasons (Kaswamila, 2012).

To facilitate implementation of its Wildlife Policy, the Tanzanian government established regulations and guidelines for the designation of WMAs (United Republic of Tanzania for Ministry of Natural Resources and Tourism, 2012). Formal implementation began in 2003, leading to the gazetting of the first pilot WMAs between 2006 and 2009. The establishment process of WMAs starts with raising awareness among local communities about the importance of wildlife conservation and the potential benefits of WMAs. Communities then elect representative members to form village level Community-Based Organizations (CBOs), which are given official authority to manage conservation activities and align them with community needs. These new CBO representatives, together with established community leaders like village chairpersons, then develop locally-customized bylaws that, together with the government Wildlife Policy and WMAs regulations, serve as the legal framework for operating each WMA (United Republic of Tanzania for Ministry of Natural Resources and Tourism, 1998, 2012). The number of WMAs at various stages of development in Tanzania increased from 17 to 37 during the period from 2003 to 2022 (Mgonja, 2023). Of these, 16 are still in various stages of establishment, such as formal application and government approval, while 21 have been gazetted and granted user rights (United Republic of Tanzania for Ministry of Natural Resources and Tourism, 2023). The performance of WMAs has been mixed; while some have achieved notable success in achieving demonstrable conservation and community development outcomes, others have struggled (Kimario et al., 2020a; Mariki, 2018; Mwakaje, 2008; United Republic of Tanzania for Ministry of Natural Resources and Tourism, 2023; USAID, 2013).

This study specifically evaluates the perceived effectiveness of the ILUMA WMA in southern Tanzania (Duggan, 2023; Duggan et al., 2024a; Duggan et al., 2024, 2024b; Kavishe et al., 2024, 2024a; Walsh, 2023; Walsh et al., 2024), focusing on its impact on local conservation, livelihoods, and community development, as viewed by various local, district, regional and national stakeholders. ILUMA began its establishment process in October 2011 and was granted user rights in May 2015, becoming the 19th WMA in Tanzania at the time. As illustrated in figure 1 is an important areas for ensuring the connectivity of three large centrally-managed protected areas nearly, namely the Nyerere National Park (NNP, previously part of the Selous Game Reserve until 2019), Kilombero Game Reserve (previously the Kilombero Game Controlled Area until 2023), and the Udzungwa Mountains National Park (CWMAC, 2019). As this relatively new WMA is surrounded by villages whose long term residents are farmers and fishers, and land pressure from more recently arrived agro-pastoralist immigrants has been mounting in the area for three decades (Brehony, 2005), it is becoming increasingly important to effectively engage local communities in conserving the area. According to a 2019 national level assessment of WMA performance, ILUMA was found to be good ecological condition, with relatively healthy wildlife populations, natural intact habitats and minimal human disturbance (CWMAC, 2019). However, recent studies indicate a major increase of illegal human activities, particularly deforestation, agriculture, livestock herding, charcoal burning, timber harvesting and even permanent settlements (Duggan et al., 2024; Duggan et al., 2024b, 2024a). Therefore, this study aims to explore stakeholder perspectives regarding the underlying reasons for these challenges and identify potential solutions to improve the effectiveness and sustainability of the ILUMA WMA going forward.

**Figure 1:**
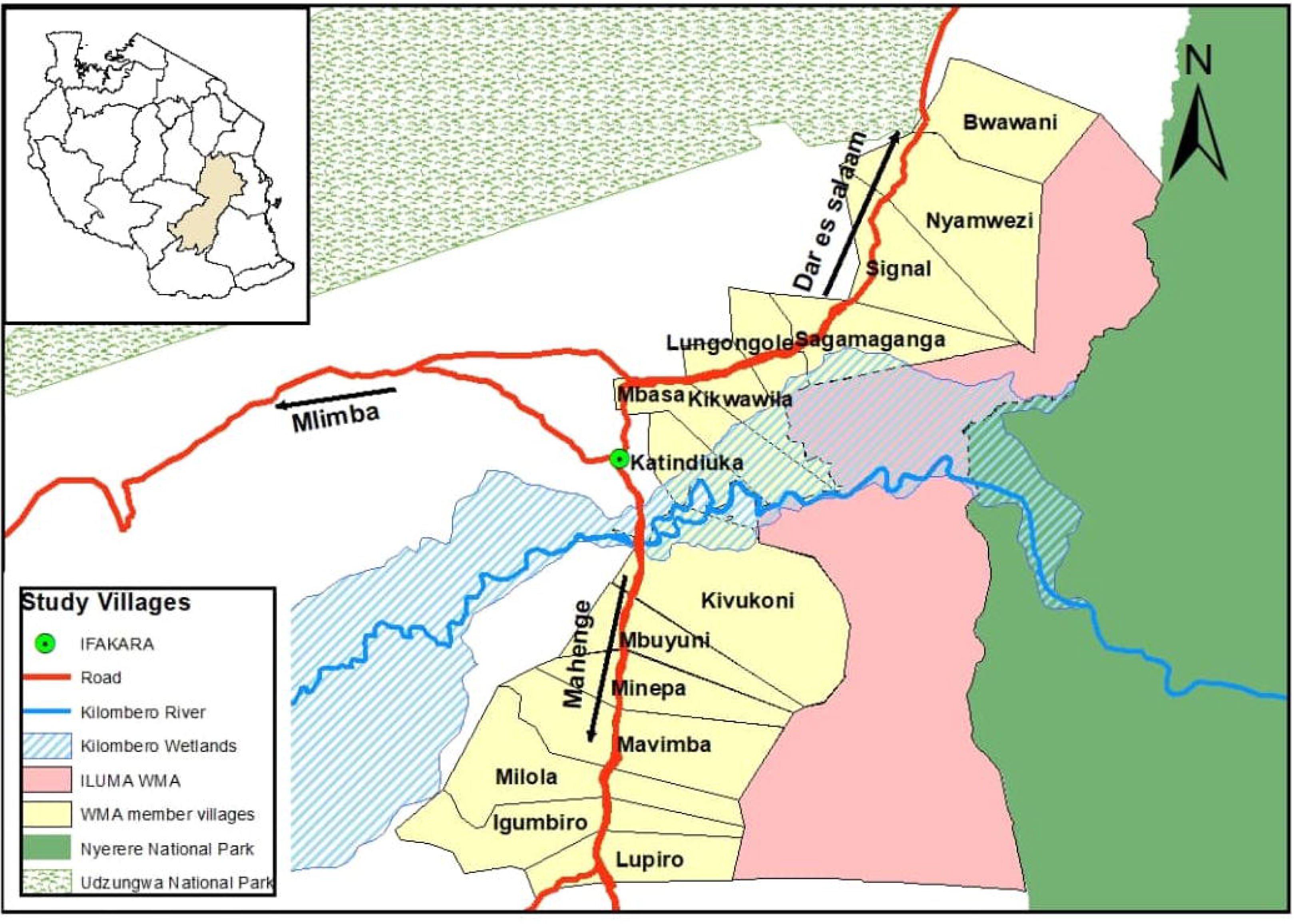
A map showing the ILUMA WMA in the east of the Kilombero Valley in southern Tanzania, with its member stakeholder villages to the west and north, from which the local community participants in this study were recruited.

## 2. METHODOLOGY

### 2.1 Study area

The ILUMA WMA in the Morogoro region of southern Tanzania (Figure 1) covers an area of 509 km^2^, spanning eastern parts of Kilombero and Ulanga districts. It is in the Kilombero valley, the largest freshwater wetland in east Africa, and serves as a buffer zone of forested habitat along the western fringe of NNP.

When the WMA was established in 2011, it comprised a group of 14 local villages, seven from Kilombero district and seven from Ulanga district (Figure 1). Later in 2014, one village (*Kiberege)* was split into three villages, namely *Bwawani*, *Nyamwezi* and *Kiberege* itself. The first of these two new villages chose to contribute their land to the WMA but the remaining *Kiberege* village chose to no longer play any part in the WMA, so it was excluded from this study. This results in a total of eight villages in the Kilombero district and 15 villages in total within the WMA.

Initially, the WMA establishment process was first supported by the Royal Belgian Government during the development and implementation of an Integrated Management Plan of Kilombero Valley Flood Plain Ramsar Site, which ended in November 2011 (Ministry of Natural Resources and Tourism, 2011). Later in 2014, the Kilombero and Lower Rufiji Wetland Ecosystem Management Project (KILORWEMP PIU, 2016) supported completion of the process in 2015. The project supported the participatory demarcation of boundaries, designation of areas for specific land use practices, formalization of the user rights, training of Village Game Scouts (community members trained to monitor and manage wildlife populations, escort visitors, prevent illegal human activities, engage local communities in conservation efforts, and address human-wildlife conflicts), capacity building of the elected WMA village representatives and facilitation of meetings with other key stakeholders.

The ILUMA WMA is rich in flora and fauna and has a range of different kinds of vegetation (Duggan et al., 2024a; Duggan et al., 2024, 2024b). Recent field surveys conducted across the WMA and some parts of the adjacent NNP demonstrated higher species richness in the intact groundwater forests, miombo woodlands and seasonal floodplain grasslands of ILUMA compared to the drier acacia savanna nearby in the NNP (Duggan, 2023; Duggan et al., 2024a; Duggan et al., 2024b). These rich environments provide favorable conditions for wide variety of wild mammals, reptiles, amphibians, birds, and fish. Regarding mammals, it is home to many large herbivores (buffalo, elephants, hippopotamuses), carnivores (lions, leopards, hyenas, wild dogs), and various antelopes (eland, reedbuck, bushbuck, red duiker, common duiker, sable antelope, suni, dik-dik, and hartebeest). Notably, the WMA hosts a significant population of puku antelope (*Cobus vardoni*) a specialized floodplain species that used to be exceptionally abundant in the Kilombero valley (Velund, 2009).

### 2.2 Research team and reflexivity

This study was conducted by the first author, who holds a Bachelor’s degree in Wildlife Management and worked as a research assistant at the time of the study. She had prior experience with qualitative research, having assisted in related studies and received training from her Masters’ degree supervisor, last author of this report and a professorial-level expert in social sciences. Throughout all the engagements with stakeholders described below, the research team maintained a neutral stance on all the subjects covered, and had no prior relationships with the participants other than their ongoing collaboration with the WMA. Additionally, the participants were not informed about the underlying hypotheses of the funded project that supported this study prior to their involvement, ensuring that their responses were as unbiased and reflective of their experiences and perspectives as possible.

### 2.3 Study design

Purposive sampling was used to select participants. The selection criteria were designed to identify individuals who hold key roles in the establishment, management and development of the WMA. Also, participants were chosen with as wide a range of different roles and experiences as possible, to ensure comprehensive representation of different perspectives from different types and levels of stakeholders within the governance and administrative structure of the WMA (Figure 2). At national level, representatives from nearby centrally-managed protected areas, under the Ministry of Natural Resources and Tourism (MNRT), Tanzania National Parks Authority (TANAPA) and Tanzania Wildlife Authority (TAWA), were recruited. These stakeholders play a role in the establishment process of the WMA and also participate in its ongoing conservation efforts through joint patrols. At regional and district level, under the oversight of the district council, a regional natural resource officer and seventeen advisory board members were recruited. These stakeholders are responsible for providing technical and administrative support to WMAs, including legal and technical advice, contract negotiations, arbitration in conflict resolution, establishment, formulation of land use plans and by-laws, and regular elections. At local level, the elected WMA village representatives, who are collectively referred as the Authorized Association (AA) empowered to manage and sustainably utilize wildlife resources on behalf of the stakeholder communities were included. Additionally, members of the village councils responsible for overseeing these AA representatives, allocating land to the WMA, and contributing to development of land use plans, were as involved as key stakeholders. Furthermore, Village Game Scouts from each WMA member village and the elected management committee for the WMA as a whole were included in the process of selecting and recruiting participants at local level.

**Figure 2:**
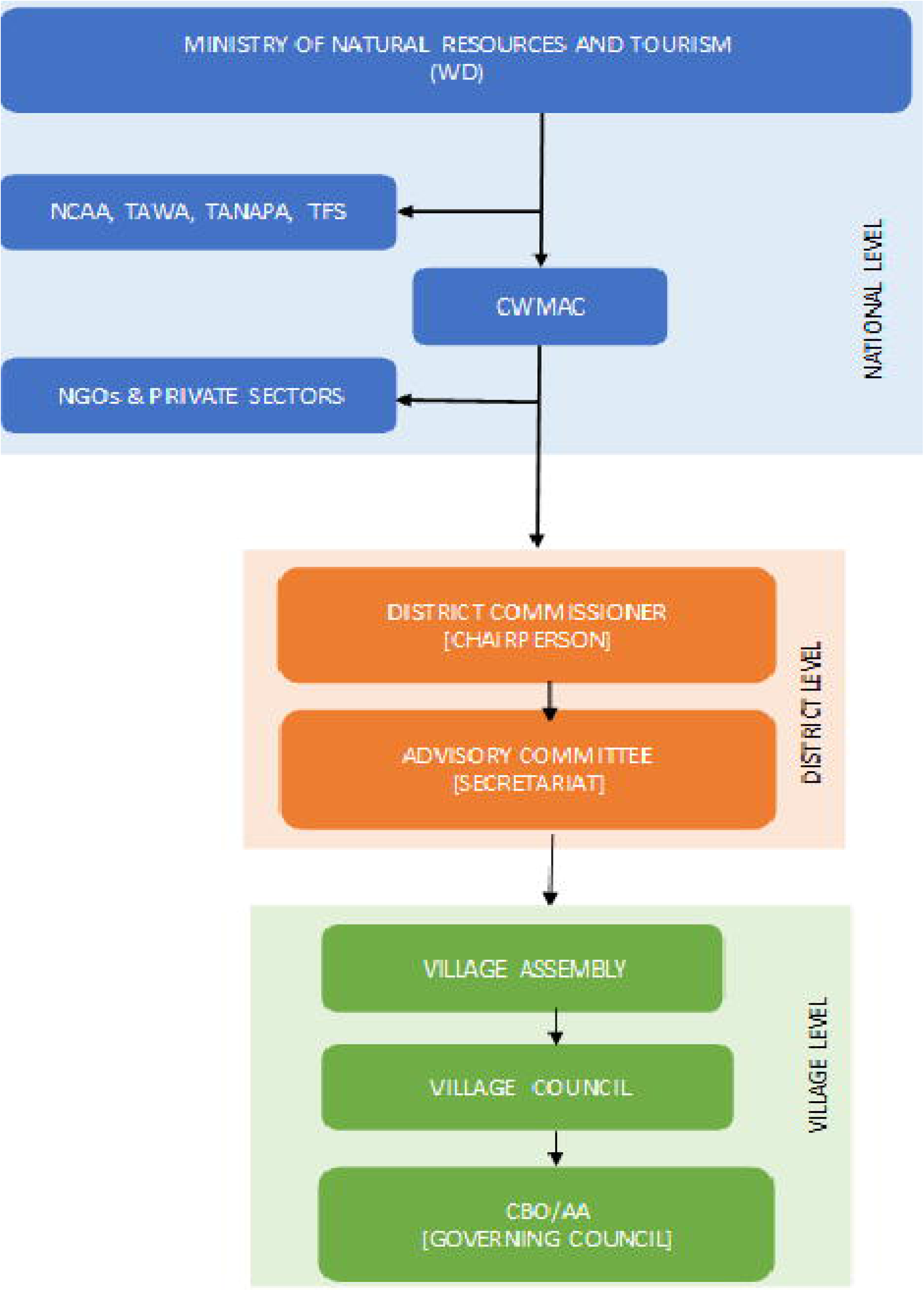
The governance and administrative structure of WMAs. DW; Wildlife Division, NGOs; Non-Government Organizations, NCAA; Ngorongoro Conservation Authority, TAWA; Tanzania Wildlife Authority, TANAPA; Tanzania National Parks, TFS; Tanzania Forestry Service, CWMAC; Community Wildlife Management Area Consortium, CBO; Community Based Conservation, AA; Authorized Association. Redrawn and adapted from (United Republic of Tanzania for Ministry of Natural Resources and Tourism, 2023).

The central methodological approach employed was face-to-face discussions in the form of in-depth interviews (IDIs) and focus group discussions (FGDs), which involved a total of 87 participants (Table 1).

**Table 1:** Numbers of various stakeholders participating in FGDs and IDIs.

| Methodology | Number of participants (and focus groups) |  |  |  |  |  |
| --- | --- | --- | --- | --- | --- | --- |
|  | Conservation<br>institutions | Regional<br>level | District<br>level | Ward<br>level | Village<br>level | WMA management<br>committee |
| IDIs | 2 | 1 | 6 | 0 | 0 | 0 |
| FGDs | 0 (0) | 0 (0) | 11 (2) | 5 (1) | 57 (12) | (5)1 |

For FGD participants, one individual was randomly selected to represent each stakeholder category in each village where multiple individuals were present, for instance the Village Game Scouts and the WMA village representative stakeholder categories. For the community leaders, WMA management committee and district advisory board stakeholder categories, a purposeful selection approach was applied to choose those most directly involved based on their roles and relevance. All recruitments of selected participants on the basis of informed consent were successful and no one declined to participate or subsequently withdrew. These formally structured sociological engagements were conducted in the village offices and workplaces of participants. We ensured that only the participants and researchers were present during these discussions, to maintain confidentiality and creating a safe space for open discussion. Demographic characteristics like gender, age, education, years at current residence, and job position or role were recorded for each participant. In addition to these prospectively collected data, complementary retrospective data sources were also used, notably recording and transcripts from a WMA stakeholder meeting held in September, 2022.

### 2.3 Data collection

#### Tool development

From personal observations, the transcripts and recordings from the public WMA stakeholders meeting served as critical background for informing the development of the survey tools (Appendix 1) and as anecdotal data sources that helped inform interpretation of the results. This meeting covered different topics regarding the overall development of the ILUMA WMA, including the benefits to the community and conservation challenges. These *a priori* experiences and information were therefore used as a basis for developing a tool for the in-depth interview (IDI) and focus group discussion (FGD) guides (Appendix 1), so that conversations with participants from this same stakeholder community flowed as freely and spontaneously as possible, while nevertheless covering all the subject matter intended.

#### Pilot study

A pilot study was conducted with a few members of the Village Game Scouts in September 2023, to test whether the questions were structured in a way that could facilitate a smooth-flowing near-spontaneous, conversation with minimal involvement of the interviewer and minimum risk of investor bias. It was observed that participants consistently narrated freely across a range of topics after the first question was asked, thereby addressing most of the remaining questions without further prompting from the investigator. These pilot study experiences indicated that the tool not only encouraged open dialogue but also fostered an environment in which participants felt comfortable sharing their insights and experiences. The clarity and relevance of the questions allowed for an easy conversation, confirming that the tool was effective in addressing the research questions, while also minimizing risk of introducing biased investigator priorities into the conversation.

#### Formal data collection

After piloting and finalizing the survey tools, the FGDs and IDIs reported herein were carried out between November and December 2023. A total of 16 FGD groups, each comprising 5 to 6 participants, were conducted with the stakeholders at village level (Village Chairperson, Village Executive Officers, elected WMA village representatives, Village Game Scouts, ward level (Ward Executive Officers), and district levels (the two most relevant WMA district advisory board members). Each FGD participant was assigned anonymized unique identifier, such as P1, P2 etc., which they were each asked to mention during the discussions, so they could be unambiguously recognized in the recordings. The IDIs were conducted with 9 individuals, specifically one senior official from NNP and another from KGR, the District Game Officers, District Commissioners, and District Executive Directors of both the Kilombero and Ulanga districts, and the Regional Natural Resource Officer for the Morogoro region.

Based on the selection criteria, which focused on stakeholders within the government and administrative structure of WMAs (Figure 2 above), FGDs were found to be well suited for engaging diverse stakeholders with similar roles within the WMA at village and district levels, to ensure comprehensive engagement with all stakeholders across 15 WMA villages, 5 wards, and the two relevant district. IDIs were considered suitable for authoritative participants at the national level, like TANAPA and TAWA, and at regional and district levels. IDIs were particularly valuable for capturing insights from higher level stakeholders, to facilitate a deeper understanding of their perspectives. Overall number of stakeholders who participated in FGDs and IDIs are shown in table 1 above.

All such engagements were conducted in *Swahili*, the common national language spoken by almost all residents of the study area. Participants were recruited through a fully documented informed consent process before the discussion began. Discussions lasted from 90 minutes up to 2 hours, during which they were digitized as audio recordings. Field notes were taken to capture contextual details and non-verbal expression to aid data analysis and interpretation. After completing the planned FGDs and IDIs, it was observed that no new themes or insights emerged from the discussions, indicating that data saturation had been reached and no additional FGDs or IDIs were necessary.

### 2. 4 Data analysis

#### Transcription and data importation

Transcriptions were done manually in the Swahili language, which involved actively listening to the audio recordings and then typing them out in a dialogue form in Microsoft *Word^®^*. Different speakers, especially in FGDs, were identified using anonymized labels like Participant 1, Participant 2, etc. Everything said by the participants was included in the transcript, even if it was outside the scope of the question asked, and written out exactly verbatim.

The data analysis for this study was conducted using *NVivo^®^* software version 20, following the thematic analysis framework. The process began with data importation of the FGD and IDI transcript files into NVivo^®^ software. The project was organized within distinct folders for data files with well labeled to ensure systematic management and accessibility throughout the analysis.

#### Case classification and coding

As a part of the data analysis, *case classification* and *coding* were conducted before the thematic analysis. Each case represents a single unit of analysis, such as an individual participant in an IDI or a specific group of participants in an FGD. Case classification means assigning attributes like demographic information to participants while case coding involves reviewing each participant’s narrative and linking their specific statements to their identities like P1, P2, IDI 2 etc. This was done to ensure easily identify which participant made each statement in the coded significant sections related to the themes during the analysis and when referencing their quotes. Two separate folders were created in NVivo^®^ software for IDIs and FGDs. Each individual IDI participant was put in the corresponding folder for IDIs and each had a unique identity based on the initials of his/her job title or institution. In the FGD folder, subfolders representing each group were named according to their respective roles within the groups (FGD Ward Executive Officers, FGD1 elected WMA village representatives, FGD2 elected WMA village representatives, FGD3 elected WMA village representatives, FGD1 Village Chairpersons, FGD2 Village Chairpersons, FGD3 Village Chairpersons, FGD1 Village game scouts, FGD2 Village game scouts, FGD3 Village game scouts, FGD1 Village Executive Officers, FGD2 Village Executive Officers FGD3 Village Executive Officers, FGD1 district advisory board members and FGD2 district advisory board members). In each subfolder, individual participants assigned a unique anonymized identity based on their participant number and stakeholder category, then followed by case coding.

#### Thematic analysis

Six steps were followed for the thematic data analysis, specifically data familiarization, initial code formulation, *theme* formulation, theme review, finalization of theme definitions and then report production (Byrne, 2022; Clarke & Braun, 2017). Whenever necessary, transcripts were then read and re-read, while supporting audio recordings were re-listened to, to ensure the accuracy and completeness of the transcripts. Notes were taken within the NVivo^®^ software during the analysis using the *annotations* and *memo* functions, to capture initial impressions and potential patterns, thereby providing a foundational understanding of the data.

Both deductive and inductive methods were applied during the development of initial codes. The deductive codes were first developed using the *discussion guide tool* (Appendix 1) and study objectives to provide a framework for organizing the data, while the inductive codes emerged through transcript review of both FGDs and IDIs. A line-by-line reading was conducted, and description-focused coding was developed by understanding the transcripts, extracting relevant information, and labeling it.

Codes were sorted and grouped based on shared relationships, and looking for commonalities between the codes, to identify the themes. Themes were then reviewed to ensure each was distinct from the others, and all coded transcripts were opened within the NVivo^®^ package to confirm whether the information coded was relevant to the respective theme. Themes were then carefully reviewed again, so that the overall message was well understood and then named appropriately.

After the analysis, a total of five themes were identified, namely *perspectives on livelihood and community development functions*, *perspectives on environmental and wildlife conservation functions*, *changes in community attitudes*, *revenue sources* and *stakeholder recommendations for the sustainability of the ILUMA WMA*.

#### Separation of observations from FGDs and IDIs views

Furthermore, the coding process was repeated to enable unambiguous separation of data from IDIs and FGDs. Using the section coded from the identified themes and sub-themes, and codes in coding references in NVivo^®^, the texts were highlighted and assigned to the respective FGDs or IDIs. This was done to obtain their frequency of occurrence, broken down by survey engagement format.

#### Varied perspective level among FGDs stakeholder categories

During the formulation of initial codes for all transcripts, variations in scale rank or levels among FGD categories regarding the first two identified themes, namely *perspectives on livelihood and community development functions and perspectives on environmental and wildlife conservation functions* was identified. Additional sub-analysis was conducted by re-reading the transcripts for each FGD category. For the first theme, deductive codes were created, namely *no information* (for those who said they did not know), *no comments* for participants who did not do so, *perceived highly beneficial, perceived moderately beneficial, perceived of little benefit* and *perception of no benefit*. For the second theme, the deductive codes were: *no information*, *no comment*, *very successful, successful, mixed* (“fifty-fifty” outcome of success and failure), *minimal success* and *unsuccessful*. This was done to better understand the varied levels of participant responses regarding WMA performance and outlook. *R-studio^®^* software were used to plot these deductive semi-quantitative coding with the *ggplot2* package.

#### Report production

A framework matrix was created in NVivo^®^ and exported to an excel file. This matrix provided a structured framework for comparing coded data across different cases. The framework consisted of rows and columns, where rows represented the cases with their demographic information (e.g., individual interview, and group participants), and columns represented the themes or codes derived from the data. Data within the framework matrix were in cell format, containing text segments coded from specific participants on the respective themes. This structured format was also used for extraction of quotes cited in the report. Additionally, different visualizations obtained from Nvivo^®^ were extracted and corresponding tables were created in Microsoft Word^®^ and PowerPoint^®^ for reporting the results.

### 2.5 Ethical consideration

This study was nested within a larger project related to both malaria vector mosquito ecology (Kavishe et al., 2024, 2024a) and CBC in the ILUMA WMA (Duggan, 2023; Duggan et al., 2024a; Duggan et al., 2024, 2024b). In securing ethical approval, we provided a thorough overview of the research objectives, methodologies, and data collection tools (Appendix 1) and management procedures. The relevant informed consent forms (Appendix 2) clearly outlined the purpose of the study while ensuring that all individuals knew they could voluntarily choose to participate in the study and withdraw at any point they wished to. Participants were given opportunity to ask questions and discuss any concerns before consenting, ensuring they were fully informed. To protect participant identities, all responses were anonymized, with anonymized unique identifiers assigned to each individual, thus ensuring that their contributions could not be linked back to them in any reports or publications. It is important to note that the information from the WMA stakeholder meeting included in this study did not require ethical approval, as this was a public meeting that preceded this study for non-research purposes and did not constitute part of the formal data collection process.

All the procedures were reviewed and approved by the Institutional Review Board of the Ifakara Health Institute (IHI/IRB/MM/035-2023), the National Institute for Medical Research (NIMRH/HQ/R.8b/Vol.I/1127) in Tanzania and Social Research and Ethics Committee (SREC) at University College Cork in Ireland. Furthermore, local government approval was provided by the relevant region and districts of the study area (Approved letters AB.175/245/01/0/55 from Morogoro region and UDC/ADM/N.10/5 VOL V/203 from Ulanga district, plus an approval letter from Kilombero district lacking a reference number).

## 3.0. RESULT

### 3.1 Demographic characteristics of study participants

A total of 87 stakeholders participated in this study (Table 1), among whom age ranged from 18 to 60 years. Two-fifths of participants (35) had primary education, while (11) had secondary education, (13) had technical education, and (28) had university education. Additionally, 33 participants were community leaders, including Village Chairpersons, Village Executive Officers, and Ward Executive Officers from the 15 WMA stakeholder villages. Among the community leader groups, 18 participants had resided at their current location for over ten years, two for 6 to 10 years, seven for 1 to 5 years, and six for less than one year.

### 3.2 Themes and sub-themes identified among stakeholder perspectives

A total of five core themes were identified, all of which were directly related to *a priori* study goals. The identified themes show how community development, natural resources management, and conservation efforts interact with each other (Figure 3).

**Figure 3:**
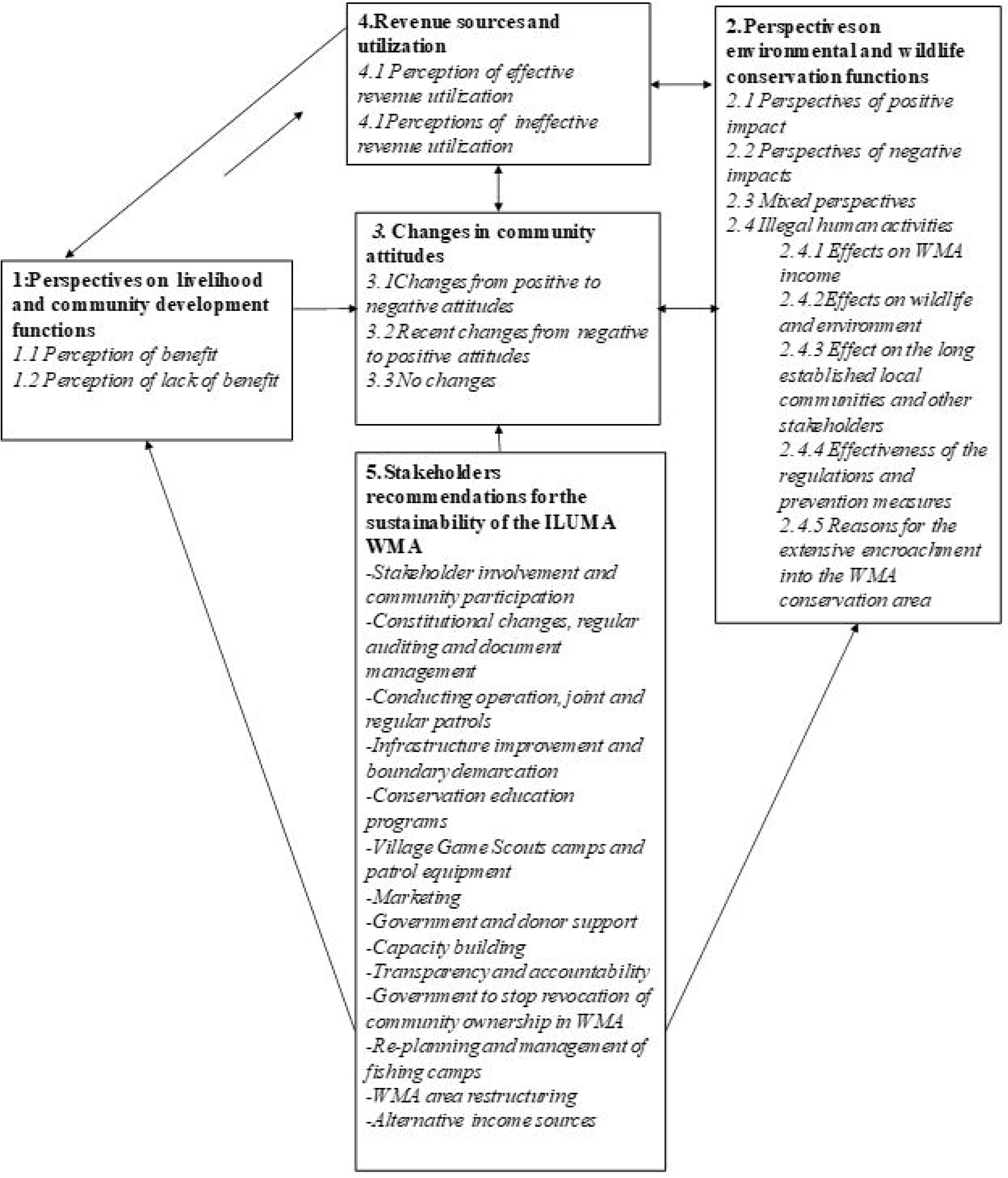
Causal relationships between the identified themes and subthemes within them.

#### 3.2.1 Perspectives on livelihood and community development functions

This study revealed mixed perspectives among the participants within groups and between the groups, regarding the WMA contributions to local livelihoods and community development. While a few participants expressed that they perceived some community benefits were accrued from the WMA, the majority emphasized lack of benefit to themselves or the community at large (Table 2). This indicates that, although the WMA was intended to improve local livelihoods and support community development, it was perceived to have not effectively achieved this goal.

**Table 2:**
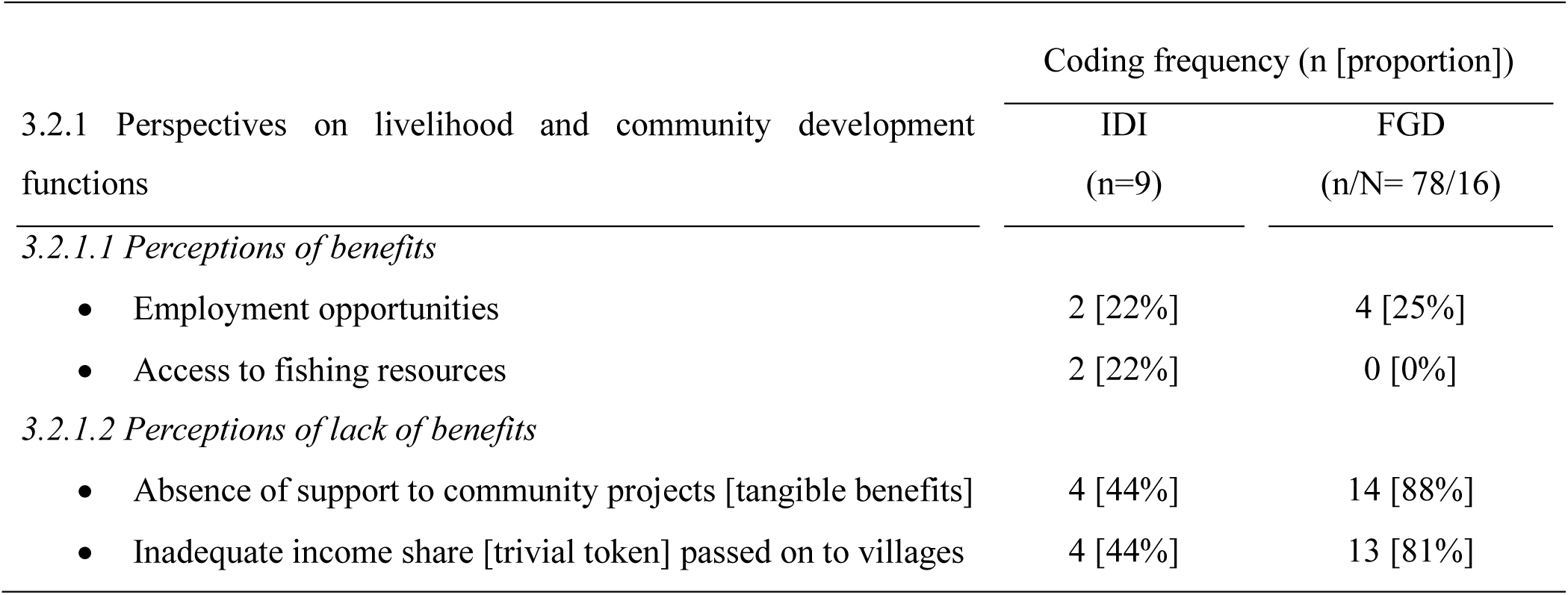
Perspectives on livelihood and community development functions of the WMA.

##### 3.2.1.1 Perception of benefits

Employment opportunities were perceived as the primary benefit realized by some community members, particularly Village Game Scouts, who are remunerated on a casual basis for escorting tourists and researchers visiting the WMA. This benefit was mentioned in just over one fifth of

IDIs and approximately one quarter FGDs (Table 2), particularly those with Village Game Scouts, WMA management committee members and district-level stakeholders:

Quote 1: *“I’ve been able to improve my daily life. I got the opportunity to work with researchers, and the money I earned helped me start a small business. Now, that business helps me provide food for my children.”* P3, FGD 6.

However, it was noted that these opportunities are accessible only to a limited number of community members who had received conservation and firearm trainings during the WMA establishment process.

Beyond casual employment as Village Game Scouts, centralized conservation institution (NNP and KGR) stakeholders participating in IDIs, mentioned access to fishing resources within the WMA area as a benefit realized by the local stakeholder communities. However, this perspective was not shared by FGD participants from the WMA villages. These participants did not mention access to fishing resources as a benefit resulting from the WMA, probably because such access existed even before the WMA was established.

##### 3.2.1.2 Perceptions of lack of benefits

The vast majority of FGD groups participants and almost half of IDI participants considered that the community has not realized satisfactory benefits from the WMA. These views were explained in terms of the lack of WMA support for community projects and trivial token income provided to the WMA villages (Table 2). All FGDs that included the village chairpersons expressed dissatisfaction with the share of income they have received from the WMA since its establishment. One village chairperson argued that the land they designated for conservation could have been more economically beneficial if used for other purposes:

Quote 2: *“The undeniable truth is that the community has not yet seen the benefits of ILUMA WMA. This is because there have been no noticeable changes in their community in terms of project support such as schools and hospitals. Therefore, the community’s development resulting from the presence of the ILUMA WMA is still lacking*.” P3, FGD 5.

Quote 3: *The conservation of this WMA has been ongoing for about 13 years, but the money we have received up to now is only Tsh 2,333,333 [$897.4] and another Tsh 99,000 [$38.1] from previous years. Can you even build a toilet with that amount? What impact does it have?… If you give the citizens even five acres to farm, they would make more than that amount of money.”* P1, FGD 7.

##### 3.2.1.3 Varying FGDs participant’s perspectives on overall WMA benefits to the community

A breakdown of perspectives among focus group participants revealed a range of views on the benefits of the WMA (Figure 4). Village chairpersons and WMA village representatives were more likely to perceive little or no benefits. Their perspectives were based on their awareness of the WMA revenue generation activities. In contrast, most Village Game Scouts expressed more positive perceptions of WMA benefit. Their involvement in WMA activities, which provide them with income as individuals understandably led to them having more generally favourable views.

**Figure 4:**
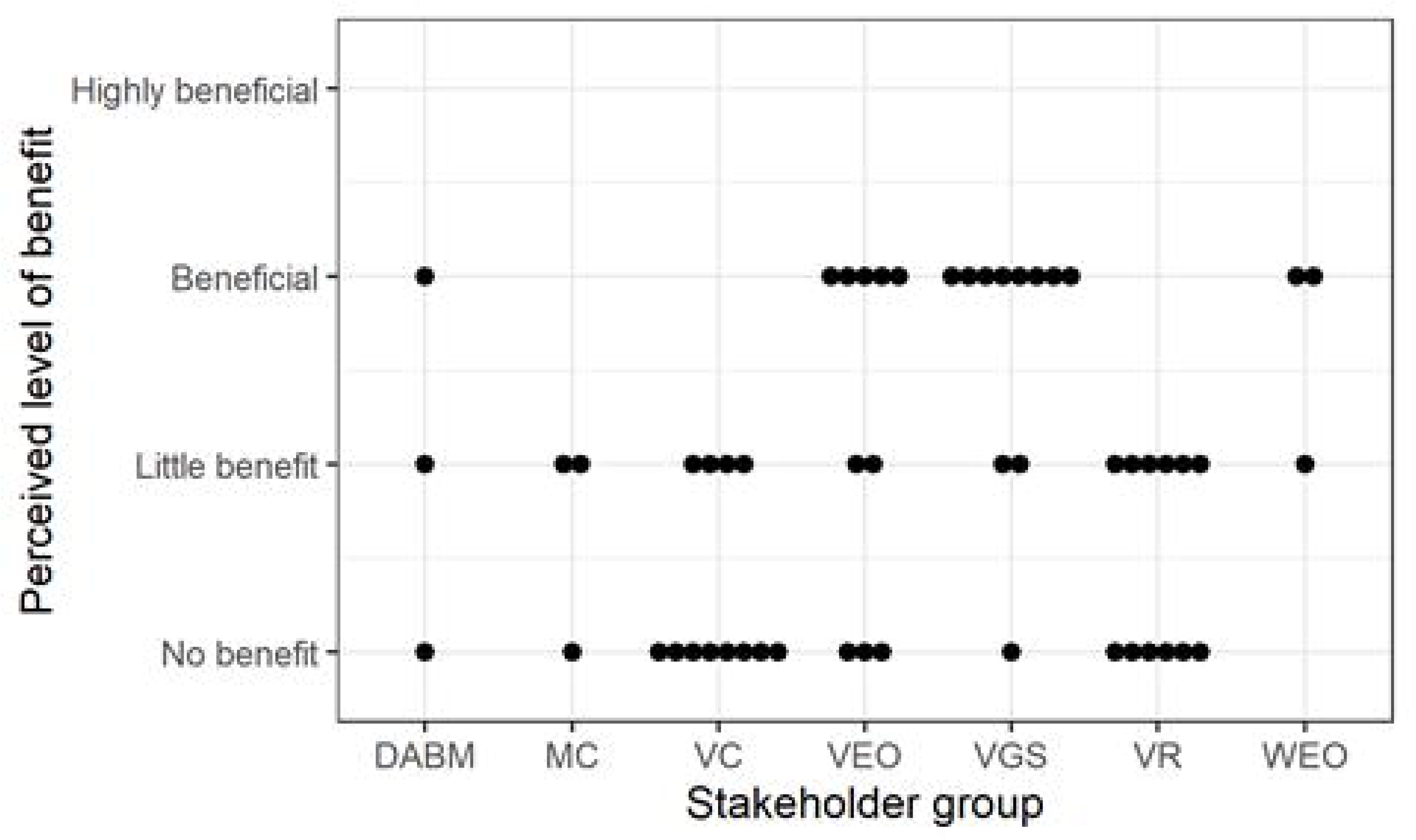
Dot plots summarizing varying responses from individual participants in FGD groups on WMA benefit to the community. DABM; District Advisory Board Members, MC; WMA Management Committee members, VC; Village Chairpersons, VEO; Village Executive Officers, VGS; Village Game Scouts, VR; WMA village representatives, WEO; Ward Executive Officers.

However, the perspectives of Village Executive Officers, Ward Executive Officers, and advisory board members from the district were mixed. Some perceive the WMA as beneficial and explained that their views were influenced by recent income distributions to villages in the year of 2023, totalling Tsh 2,000,000 [$769.2] per village. Others lacked information about the WMA benefits, but it should be noted that most of these individuals did not originate from the area and were government employees who were frequently relocated.

In the FGD with Ward Executive Officers, one reported that WMA management committee members and village chairpersons do not transparently disclose revenue information or involve the community in decision-making:

Quote 4: *“In the village meetings, the chairman does not announce and let the community know if WMA has given them the revenues to support their projects. Even WMA management committee, they don’t follow up with the revenues they brought to the villages. I once visited one of my villages and I didn’t find any information regarding the income received from this WMA.”* P2, FGD 13.

As a result, the accrued income is usually known only to the village chairpersons. However, during discussions with village chairpersons, one explained that the amount of money received was too small to justify meaningful community decisions. Consequently, the funds were used for maintaining and repairing the village offices and school classrooms.

#### 3.2.2 Perspectives on environmental and wildlife conservation functions

Perspectives regarding the conservation functions of the WMA were mixed within and between stakeholder groups. Some views were positive, recognizing the WMA contributions to conservation efforts, while others expressed concerns about negative impacts on the environment resulting from its establishment (Table 3). These varied perspectives indicate that the WMA has experienced both successes and failures in achieving its conservation goal.

**Table 3:** Perspectives on environmental and wildlife conservation functions.

|  | Coding frequency<br>(n [Proportion]) |  |
| --- | --- | --- |
|  | IDI<br>(n=9) | FGD<br>(n/N =78/16) |
| 3.2.2 Environmental and wildlife conservation functions |  |  |
| 3.2.2.1 *Perspectives of positive impact* |  |  |
| • Reduce poaching | 7 [78%] | 11 [69%] |
| • Contribute to forest protection | 5 [ 56%] | 4 [ 25%] |
| 3.2.2.2 *Perspectives of Negative impact* |  |  |
| • Increase of human encroachments | 3 [ 33%] | 13 [81%] |
| 3.2.2.3 *Mixture perspectives* |  |  |
| • “Fifty-fifty” mixture of success and failure | 1 [11%] | 7 [44%] |

##### 3.2.2.1 Perspectives of positive impact

Most positive perspectives were based on a reduction in wildlife poaching following the establishment of the WMA. This reduction is attributed to increased community involvement in wildlife conservation, as noted in almost all IDIs and in about two third of FGDs (Table 3):

Quote 5: *We used to have so many cases of wildlife being killed in most of this village’s lands adjacent to our protected areas. However, after the establishment of this WMA it is very rare to hear about such issues. This means that our WMA helps conserve the wildlife.”* IDI 3.

It should be noted, however, that the reduction in poaching might also be partly due to the Tanzanian government’s broader anti-poaching efforts at the time.

Additionally, more than half of IDIs and one-quarter of FGDs perceived that the WMA plays a crucial role in conservation by acting as a buffer zone for NNP. They considered that its establishment had contributed to the protection of wildlife and their habitats, all of which might otherwise have been disturbed by human activities. For instance, discussions with village chairpersons, Village Game Scouts and WMA village representatives from Kilombero district reported that forests on their village lands, previously threatened by human activities, had recovered following upon the establishment of the WMA:

Quote 6: *“In some other villages like our village, we have a big forest. Before the establishment of the ILUMA WMA, resources used were unregulated, and people used to enter and do any activity they wanted to do. But after the forest was set as a conservation area, the forest got much more protection. Also, there were areas where there was a lot of illegal fishing using nets with small holes. After becoming part of a WMA, such illegal activities decreased. Pastoralists are still in there but stay cautiously, knowing it’s a conserved area.”* P3, FGD 10.

This recovery resulted from improved protection by the local community themselves and new regulations implemented after the WMA was established. However, human encroachment was considered to remain a limitation, requiring ongoing efforts to address the pressures on these lands and ensure sustainable management.

##### 3.2.2.2 Perspectives of negative impact

On the other hand, in over three-quarter of FGDs and one third of IDIs, participants expressed the view that the WMA had not met its conservation goals. The chairpersons in Ulanga district villages explained that most of the village land had a better conservation status in the past, before the WMA was established, compared to now. Large parts of what are now WMA village lands were rich in wildlife and had been used for tourist hunting activities (Brehony, 2005) but were now experiencing increased human encroachment into the areas designated for conservation.

Quote 7: *“If the current WMA conservation status had been the situation when the idea of the WMA was proposed, we wouldn’t have agreed to set our village land aside for conservation. Even those who supported the WMA establishment would not have wanted those areas included, because now the area is encroached by invaders who are living and farming there, but previously before WMA was established no one was living there.”* P4, FGD 7.

Quote 8: *“I can’t say ILUMA is doing well in conservation because some WMA areas have turned into villages where people even run businesses. (Laugh)…I think the Tanzania Revenue Authority should collect taxes from these activities. It’s surprising to see people setting up small shops and selling goods in the conservation area.”* P4, FGD 5.

Consistent with the direct observational surveys of human, livestock and wildlife activities recently carried out across the WMA (Duggan, 2023; Duggan et al., 2024a; Duggan et al., 2024), one stakeholder from NNP and an ILUMA village representative from Ulanga district noted that some parts of the WMA now resemble small villages because of the many new settlements established by *waSukuma* agro-pastoralist immigrants from western Tanzania. The negative sentiments expressed regarding this relatively recently arrived ethnic group are consistent with reports dating back to over two decades ago (Brehony, 2005) and are emphasized by the frequent use of the Swahili word *wavamizi*, meaning *invaders*, which featured 45 times across all transcripts.

It was, however notable that participants from Kilombero district did not share this consistently negative view of the *waSukuma* community settlements; however, they did express concern about livestock herding during the dry season. These differing views may well arise from the distinctive ecology and settlement patterns across the WMA, with the miombo woodlands to the south of the Kilombero river being for more vulnerable to encroachment than either the groundwater forest along its south bank that is occupied by authorized, regulated resident fishing communities who act as custodians of the local ecosystem (Duggan et al., 2024b), or the floodplain grassland to the north of it that cannot be settled because it is regularly inundated.

##### 3.2.2.3 Mixed Perspectives

Almost half of FGD groups and one individual IDI, expressed mixed views on the effectiveness of the WMA conservation achievement, which some participants summarized as a “fifty-fifty” outcome:

Quote 9: *“Myself, I would say on the scale of hundred, we are fifty-fifty, with half destruction and half conserved. We still have wild animals present in some of our WMA area”* P2, FGD 14.

One district stakeholder, whose duties include overseeing the WMA, and who has resided locally for seven years, explained that he has been told that there were minimal human activities encroaching on the land when the WMA was first established. He/she indicated that between 2018 and 2020, the area experienced heavy encroachment, resulting in extensive environmental destruction, an account that was verified by recent field surveys (Duggan, 2023; Duggan et al., 2024a; Duggan et al., 2024). Lastly, he/she reported that some improvements have been recently observed, with a decrease in encroachment in certain areas that were previously heavily affected:

Quote 10: “*There is an area called Mdaba Valley, five to six years ago there were hectares cultivated in that entire area up to the Kilombero River. But if you go there now, that area has not been cultivated for almost two years, and now its natural vegetation has recovered. Even wild animals have started getting back there, especially in the dry season.”* IDI 8.

Having said that, perceptions of conservation effectiveness varied considerably across different stakeholder groups, and it was notable that no participant considered the WMA to have been very successful (Figure 5). Many FGD participants perceived that the WMA had either achieved minimal success or had been successful. However, two participants indicated that the WMA had been completely unsuccessful regarding conservation, while seven others considered the outcome thus far to be “fifty-fifty”, meaning the area is partially protected and partially encroached. The varied perspectives were, of course, understandably influenced by their different roles (Figure 5). The views of Village Game Scout participants, for example, could be readily shaped by their involvement in patrol activities, through which they readily observed changes over time across the entire protected area of the WMA.

**Figure 5:**
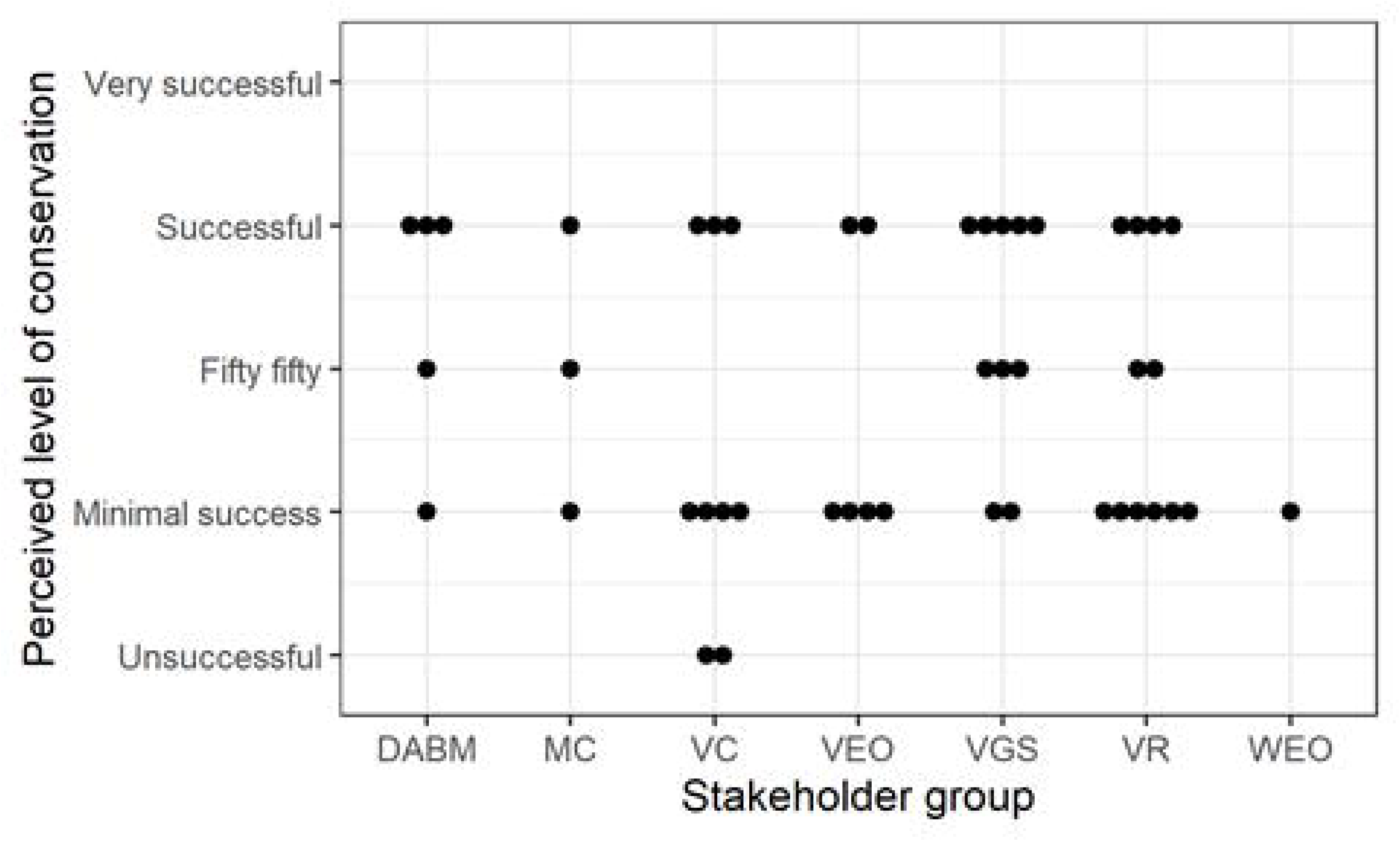
Dot plots summarizing varying responses from individual participants in FGD groups on WMA conservation effectiveness. DABM; District Advisory Board Members, MC; Members of the WMA Management Committee, VC; Village Chairpersons, VEO; Village Executive Officers, VGS; Village Game Scouts, VR; WMA Village Representatives, WEO; Ward Executive Officers.

WMA management committee members and Ward Executive Officers expressed few concerns, though their small numbers limit the certainty of this evidence. Likewise, the relatively few participating district advisory board members expressed largely positive or mixed perspectives, perhaps because they are government employees, many of whom have not resided in the area for a long time. Interestingly, other participants from several Village Executive Officers, Ward Executive Officers and one district advisory board member exhibited obvious lack of knowledge about the WMA, so their uncertainties may have contributed to the variation perspectives expressed by these stakeholder categories.

##### 3.2.2.4 Illegal human activities

Consistent with recent observational field surveys (Duggan, 2023; Duggan et al., 2024), a wide range of illegal human activities within the WMA area were mentioned in all FGDs and almost all IDIs. The most frequently mentioned activities were livestock herding, agriculture, charcoal burning, settlement, timber harvesting and fishing using nets with small holes. Other reported activities were firewood collection, cutting building poles, canoe making, grass cutting, mining, sand collection and creating traditional alcohol production (Figure 6).

**Figure 6:**
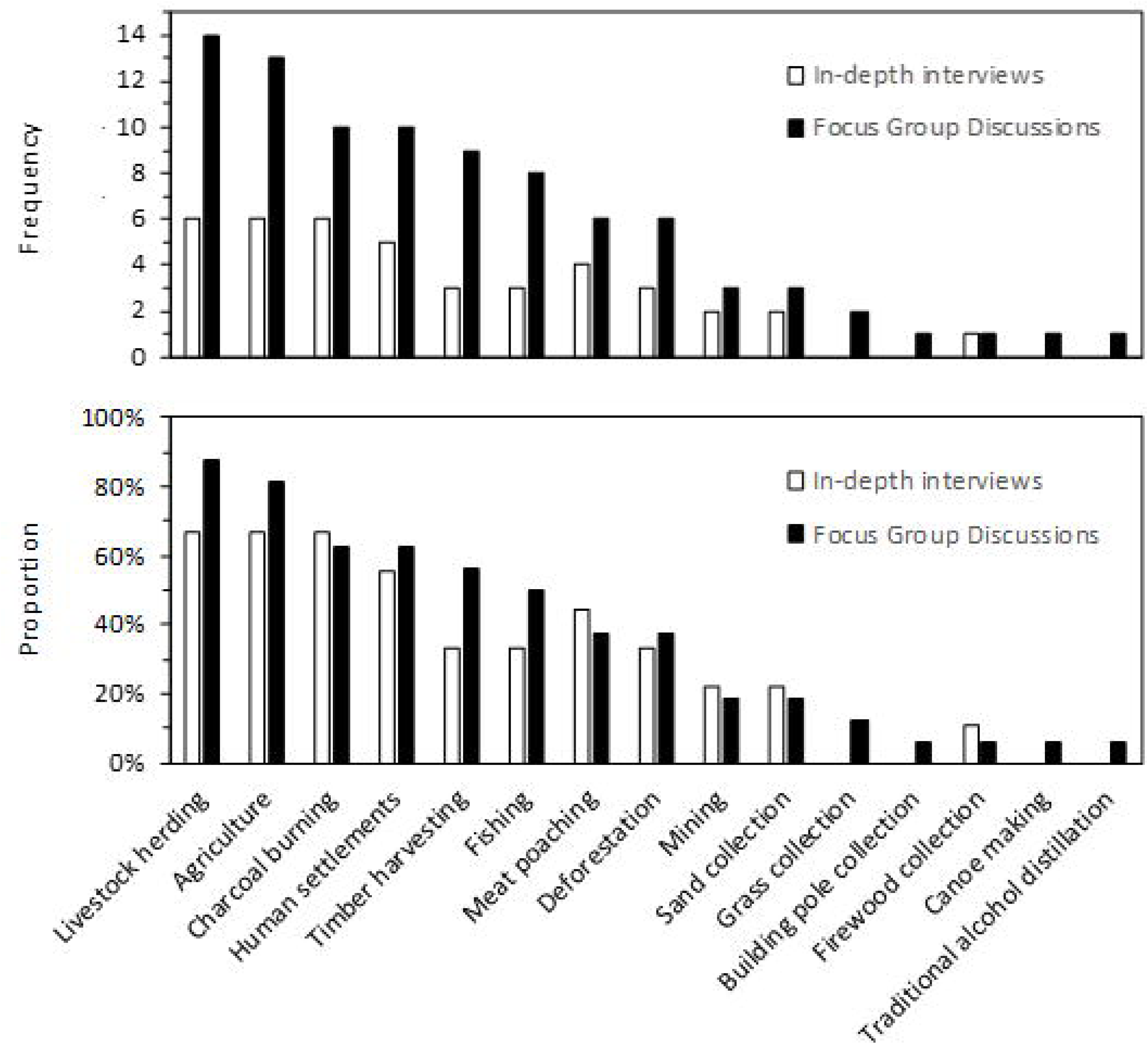
Frequency and proportional frequency with which various illegal human activities were mentioned in IDI (9), FGDs (16) by study participants (87).

Quote 11: *“The WMA area has been like a grandmother’s farm that anyone can use [freely]. There are livestock herding, farming, illegal fishing, charcoal burning, illegal poaching, sand collections, illegal mining, timber harvesting. Those are the challenges. Others are settlements. People are even hiding there making local brews along the rivers.”* P4, FGD 7.

Factors considered relevant driving these illegal activities included land fertility, the presence of suitable trees for charcoal and timber, demand for bush meat and need for income among surrounding communities. District Game Officers, Village Game Scouts and WMA management committee, all reported that most illegal activities, such as livestock keeping and agriculture, were carried out by agro-pastoralists who had recently immigrated from western Tanzania and established settlements adjacent to and even inside the conservation area. Conversely, charcoal burning, timber harvesting, illegal fishing, and poaching were mostly considered to be conducted by the long-established communities in the area, most of whom have lived there for well over a century (Larson, 1976):

Quote 12: *“Approximately ninety percent of the encroachers are not the long resident local people who set aside their land for conservation. The long-term residents have care for their conserved area but mostly other people from far away in Shinyanga region are the ones who have established settlements in there and do farming. You can rarely find locals have settle inside there. However, most of them* [who do enter the WMA] *are charcoal burners or timber harvesters who normally do not establish settlements.”* IDI 3.

Further findings reveal that illegal activities not only affect the conservation of the environment and wildlife within the WMA but also negatively impact the local community, the WMA as an institution and several other associated stakeholders (Table 4).

**Table 4:** Perspectives on the effect of illegal human activities.

|  | Coding frequency<br>(n [proportion]) |  |
| --- | --- | --- |
|  | IDI | FGD |
|  | (n=9) | (n/N=78/16) |
| *3.2.2.4 Illegal human activities* |  |  |
| *3.2.2.4.1 Effects on WMA income* |  |  |
| • Investors lost interest | 2 [22%] | 9 [56%] |
| • Unsatisfied tourists | 2 [22%] | 6 [38%] |
| *3.2.2.4.2 Effects on wildlife and environment* |  |  |
| • Shifting of animals away from WMA | 2 [22%] | 9 [56%] |
| • Destruction of water bodies and wild habitat | 1 [11%] | 4 [25%] |
| • Increases threats to nearby parks and reserves | 1 [11%] | 0 [0%] |
| *3.2.2.4.3 Effect on the long-established local communities and other stakeholders* |  |  |
| • Negatively affects activities of research collaborators | 0 [0%] | 4 [25%] |
| • Increase of crop destruction by animals | 0 [0%] | 3 [19%] |
| • Loss of cultural needs from natural forests | 0 [0%] | 1 [6%] |
| • Fear for safety of visiting tourists | 1 [11%] | 1 [6%] |

###### 3.2.2.4.1 Effects on WMA income

More than half of FGD groups and about one-fifth of IDIs revealed the widely held view that potential investors have lost interest in the WMA due to the extensive human disturbance that had been clearly documented across approximately two thirds of the WMA (Duggan, 2023; Duggan et al., 2024a). Additionally, District Game Officers and Village Game Scouts reported that tourists were dissatisfied with their visits.

Quote 13: “*Sometimes our tourists get dissatisfied with the way our conservation status changes. We used to walk short distances and see a lot of wild animals but today it’s no longer like that. We walk for so long to make our tourists satisfied with what they wish to see. It is really challenging. Imagine you showed a tourist a good tree and he or she likes it, then the next time she or he comes back again and wants to see the same tree again and it’s no longer there. It is really disappointing.”* P2, FGD.

Moreover, at a public stakeholders meeting, a prospective investor expressed disappointment with the extensive encroachment within the WMA.

Quote 14: *“Sometimes it’s challenging, and it is even disappointing. We decided to support the patrols in the hunting block I am expecting to invest in, but we usually meet with the charcoal burners and so many groups of cattle inside the WMA, so my client got dissatisfied. Sometimes I had to get them in during the night, so that they won’t see cattle groups on the way to the block.”* Stakeholders public meeting.

###### 3.2.2.4.2 Effects on wildlife and environment

Participants from a variety groups reported that illegal human activities have had several negative impacts on wildlife and the environment. Many explained that animals were moving away from the WMA. Additionally, water bodies and wildlife habitats had been damaged due to livestock herding and agricultural activities, as noted by some Village Game Scouts. One District Game Officer expressed a concern about the issue of increased threats to adjacent centrally protected areas, especially NNP. Indeed, he reported that the WMA is used as a staging post into the adjacent park by poachers who enter and hunt inside it:

Quote 15: *“Meat poachers use ILUMA as a pathway to get into the NNP for hunting, then after they get back to smoke their meat in the WMA because they are afraid to spend more time in the NNP”* IDI 8.

He also added that livestock herding inside the WMA sometimes extends into the park:

Quote 16: *“Even if you walk through some areas in the WMA you can see the livestock tracks crossing the boundary and entering to the NNP”.* IDI 8.

###### 3.2.2.4.3 Effect on long established local communities and other stakeholders

The perspectives shared by some District Advisory Board Members and Village Game Scouts indicate that illegal activities had negatively impacted research activities that were conducted within the WMA. Furthermore, participants from the northernmost villages reported an increase in crop destruction by animals, elephants in particular, which negatively impacted many households and the community as whole. Interestingly, one of the FGDs debated whether this increase in crop damage was due to the conservation successes of the WMA or its failures, with the latter possibly resulting in habitat destruction that pushed these animals into village farms:

Quote 17 “*I do not think the increase of elephant coming in our settlement is because ILUMA is performing well. It might be because these animals are getting confused with their environment. Maybe elephants are getting lost. Just from nowhere they find themselves boom…at the wall of people’s house*.” P5 FGD 5.

Additionally, one Village Chairperson expressed frustration that community elders can no longer perform their traditional rituals, such as rainmaking ceremonies in their village forests, due to increased deforestation:

Quote 18: “*We used to ask for rainfall from dense forests, but now we can no longer do it because the big trees we used are now all cleared*”. P5 FGD 10.

Lastly, two participants expressed concerns that illegal human activities inside the WMA might cause fear among visitors.

Quote 19: *“Sometimes it gives us difficulties to explain to our visitors when they see the encroachments within our WMA. They always ask what is this, or who is that, and why he/she is here while it’s a conservation area? It might even make them feel unsafety when they see people around.* P3, FGD 3.

###### 3.2.2.4.4 Effectiveness of the regulations and prevention measures against illegal activities

Discussions with WMA village representatives, Village Game Scouts, WMA management committee, and District Game Officers revealed that fines are the primary means used to punish illegal activities, particularly cattle herding, while patrols serve as the main preventive measure. However, most participants viewed these measures as ineffective (Table 5). They explained that, when cattle are caught in the WMA, the owners often pay fines to retrieve their cattle, and many are willing to do so. Indeed, cattle apprehended by the VGS frequently return to the WMA even after multiple cycles of fines. Additionally, a District Game Officer explained that patrols are conducted only occasionally and faced transport limitations. Village Game Scouts explained that they usually travel to the WMA by motorbike and conduct patrols on foot, making it difficult to cover large areas effectively.

**Table 5:** Frequency of mention about the effectiveness of the regulation measure against illegal activities from the transcript review of both IDIs and FGDs.

|  | Coding frequency<br>(n [proportion]) |
| --- | --- |
| *3.2.2.4.4 Effectiveness of the regulations and prevention measures against illegal activities* | Participants<br>(N= 87) |
| • Perceptions of effectiveness<br>• Perceptions of ineffectiveness<br>• Perceptions of a mixed “Fifty-fifty” picture | 1[ 1%]<br>22[28%]<br>3 [3%] |

Quote 20*: “We are struggling with the patrols, which I can say are ineffective because we have no working tools. We only have very few firearms that we got funded by the researchers* [Investigators for the research project reported herein and elsewhere (Duggan et al., 2024; Kavishe et al., 2024; Walsh et al., 2024)]. *We do not have vehicles that could make it easy to conduct regular and efficient patrols, so you may find that within five days of patrols you cannot even cover a quarter of the area. You find other areas remain with no patrols at all”* IDI 3.

Quote 21*: “I have a Sukuma friend, he told me if someone succeeds well at staying in the conservation area for just two years without being caught, if they get caught in the third year, they can pay any amount of fine, even twenty million Tanzanian shillings [$7,692]. So, we need to change this rule, we can see these fines are a lot of money for us but for them means nothing, This Sukuma tribe, when their livestock get caught and pay fines, they take this easily as a normal challenge in their life. He even told me that they are happy with getting trouble with their cattle’s because they have a cultural belief that the more they get in trouble the more their cattle produces. Using fines, we will never be able to control these people.”* P4, FGD 5.

On the other hand, a few participants viewed these measures as somewhat effective. They explained that fines deterred the encroachers from entering the conservation area.

Quote 22*: “But at least we are not just all quiet and let them encroach as they want. We do make the encroachers pay fines when we catch them in our WMA with cattle. So, I think, somehow, they get deterred from entering the WMA.”* P4, FGD 11.

###### 3.2.2.4.5 Reasons for the extensive encroachment into the WMA conservation area

IDIs and FGDs mentioned several reasons for the increasing frequency of illegal activities within the WMA. Most of FGD participants pointed to political influences, corruption, and management challenges. In contrast, IDIs more frequently mentioned the upgrade of nearby protected areas (KGR and NNP), while financial constraints were noted in more than half of both IDIs and FGDs. Other reasons included a lack of conservation education, inadequate support from the district government, insufficient patrol equipment, irregular patrols, and insufficient attention to charcoal burners by Tanzania Forest Services (Table 6).

**Table 6:**
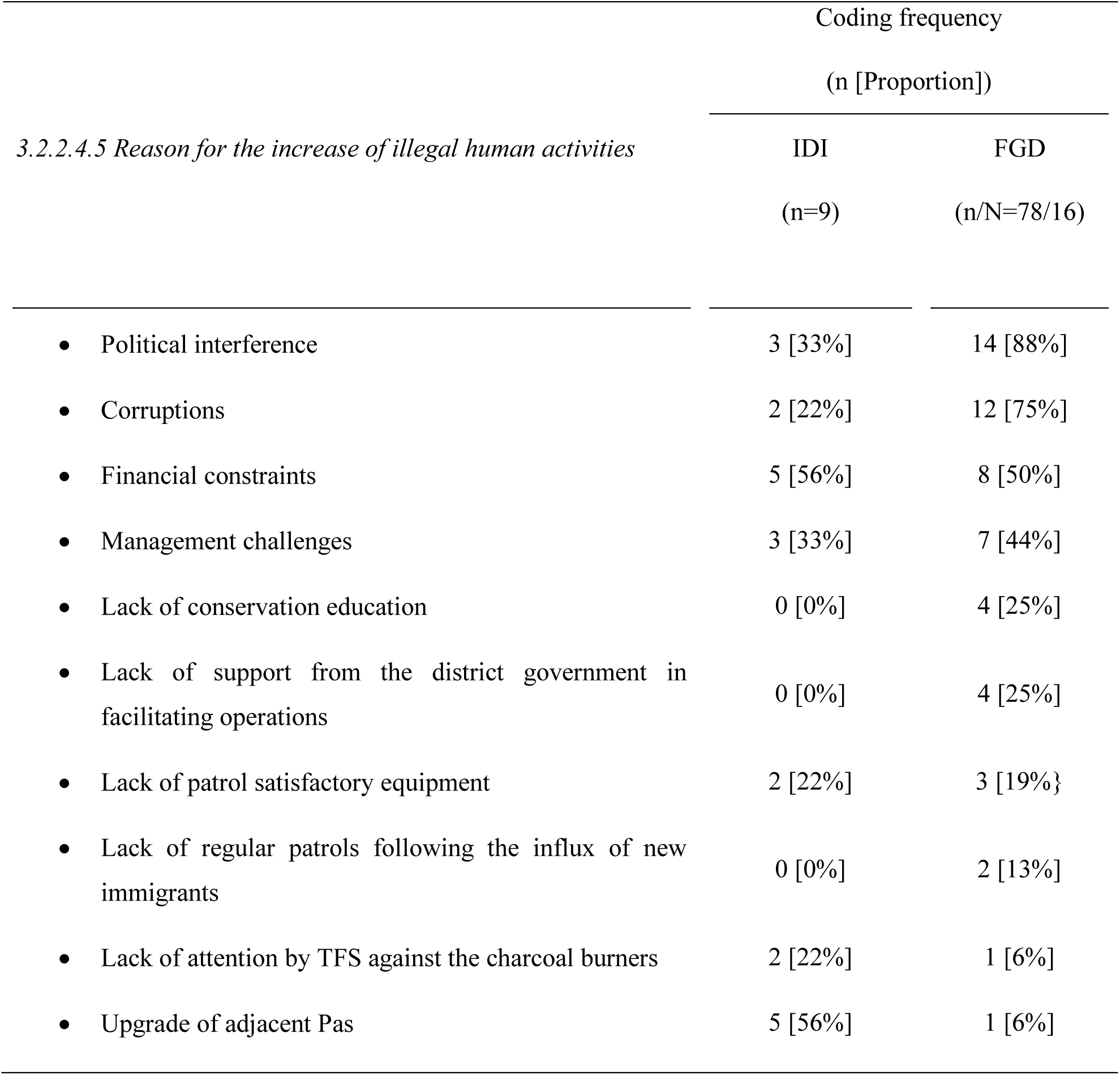
Perspectives on reasons for the increase of illegal human activities within the WMA.

###### Political interference

Participants from almost every FGD and one-quarter of IDIs reported that the elected community representatives, for instance the councillors and village chairpersons, take advantage of the conservation area by convincing people that they will be able to distribute the land for farming and livestock keeping if they vote for them. This increases encroachment pressure as people believe they will eventually own the land. However, this issue was less frequently noted in the discussions with village chairpersons FGD, which is perhaps unsurprising as they are subject of such narratives.

Additionally, WMA management committee and some of Village Game Scouts explained that the arrived new immigrants have even been informally allocated some areas, which has increased their confidence to encroach, believing they have the political backing to defend them. Furthermore, they reported that even most of the recent patrols were disrupted because of the elected community representatives reporting false information to the district government:

Quote 23*: “Sometimes the elected representatives of community leaders interfered. Once operations to evict people from the conservation area are to be conducted, suddenly you hear them reporting false information to the district government that operations are causing people injuries and other rumours. At the end you find the operations becoming difficult to carry out smoothly due to the interference. So political [local level] interference is the big problem here.* IDI 3.

Quote 24: *“As leaders, we’re doing our best to ensure good conservation of our WMA. Sometimes it’s challenging because when the cows are caught inside the WMA, the village chairpersons and councillors are the first to come and try to defend [the owners].”* Stakeholders public meeting.

For instance, they explained that they once conducted joint work patrols with rangers from TFS and TAWA, and two days later they received a notice from the District Commissioner ordering them to stop. Sometimes they were ordered to let encroachers harvest their crops inside the conservation area with a promise that they can remove them from the area afterward but instead this situation has been allowed to persist for years.

###### Corruption

The regional stakeholder and both District Game Officers in IDIs, as well as three-quarter of all FGD groups, explained that one of the reasons behind the situation was corruption within the former WMA management committee that had previously been in power. A District Game Officer reported that these leaders had a habit of taking money from people and allowing them to enter the conservation area. He explained further, saying that, because of that, the village chairpersons responsible for the land felt it was only benefiting a few people. Some of them began unofficially allocating it to new immigrants in exchange for covert cash payments.

Quote 25*: “The encroachment increased in the WMA area, as it was rumoured and observed that the former management committee of the WMA contributed to such ongoing destruction in the WMA. Village chairpersons started allocating part of the conservation area to the new immigrants illegally after they observed they were not benefiting from their land but only the elected management committee. So, everyone started to find a way that they could benefit personally from the WMA land.”* IDI 3.

It was also reported by Village Game Scout from one particular village and one former WMA village representative at the stakeholder meeting, that people are deceived regarding the WMA boundaries. One Village Game Scout explained that when they asked people who have established settlements inside the conservation area on how they got the land, some of them claimed to have bought it, and most of them only became aware that they were inside the conservation area after being there for a while:

Quote 26: *“Some individuals deceive and receive money from the encroachers, then allow them to enter the WMA by lying about the boundaries. They even assure them that they will be safe staying there. Unfortunately, the Sukuma tribe has a habit of calling their fellows when they reach the land where they think they can graze their livestock.”* Stakeholders public meeting.

These participants also explained that the same practice of deceit also occurs among these encroachers themselves. When they realize they have settled inside the conservation area, they seek out their uninformed fellows and sell the same piece of WMA land to them. Consequently, day by day, the area becomes increasingly encroached upon. As a result, it becomes difficult for these encroachers to easily accept leaving an area they have been sold illegitimately, even if they had already learned that they had done so illegally within the conservation area.

###### Financial constraints

Half of FGDs and more than half of IDIs reported that the WMA lacks sustainable income-generating activities. These financial constraints impact the ability to cover essential costs like payments to Village Game Scouts or providing food during patrols, much less pass on any surplus to the stakeholder communities. As a result, the WMA struggles to effectively address challenges within the area:

Quote 27*: “From what I know, ILUMA doesn’t have enough revenue to incur the required conservation costs. They do not have the financial power to operate patrols for controlling illegal activities, and conservation needs a lot of money. We cannot blame them that they are not working-it’s because they have no money to cover the conservation costs.”* IDI 1.

###### Management challenges

Almost half of FGDs and one third of IDIs revealed that limited management capacity at the WMA also facilitated illegal activities within the conservation area. These limitations include poor management practices, a lack of effective leadership skills, inadequate decision-making processes and insufficient coordination among stakeholders:

Quote 28: “*The WMA has a management committee, but I don’t think if they are skilled enough to operate the WMA. That is why are now about to fail to conserve the area.”* IDI 2.

For example, one village chairperson complained that the elected WMA management committee are not effectively involving the village chairpersons or community members in management decisions and activities. As a result, there is a perception that the WMA now belongs to elected WMA committee members, rather than being a shared responsibility with the community. It was considered that this lack of community involvement may contribute to the rise in illegal activities, as the community feels disconnected from the management and their conservation efforts.

Quote 29: “*The problem is that those who we selected to act as the management committee, they think have power over the control of all resources by themselves and forget about the community and other stakeholders. They must have known who our resources belong to because now it has been like they own our WMA.”* P5, FGD 10.

###### Lack of conservation education

FGD discussions with community leader groups indicated a lack of conservation education among local communities, which they perceived had led to a lack of awareness among community members regarding the importance of conserving the WMA. One Village Executive Officer who had resided in the area for more than ten years, explained that the community was only informed about conservation during the initial establishment of the WMA, and no educational programs had been implemented since then within the local community. Another expressed concerns about lack of institutional memory because the majority of individuals engaged in illegal activities inside the WMA are teenagers. She/he perceived that this group is largely unaware of conservation issues, because they were too young and had never had the opportunity to learn about conservation activities or how the WMA functions.

Quote 30: “*My concern is about conservation education. For instance, when we hold a meeting at my village, most of the population who question this WMA are youths. This may be because they know nothing about it and were not adults at the time the WMA was established. This group are the one who most frequently enter in the WMA to conduct illegal activities. Down here in the community, people are unaware about the conservation activities. If you tell them they are not allowed to enter in the WMA, they wonder and ask why?”* P1, FGD 8.

###### Lack of support from the district government

One quarter of FGD groups reported a lack of support from the relevant district governments as a limitation to effective conservation efforts. The WMA management committee revealed that they often receive little support or no response when seeking approval from the District Commissioner regarding operations to evict people from the conservation area, a situation that has persisted for a long time.

Quote 31: *“Our leaders from the district level are the ones who stop us from reaching our conservation goal because they do not give us support we need. We need their approval for operations to evict people from our area but it has been long time now. If we ask for it, we don’t see their effort on that.*” P2, FGD 14.

Additionally, during discussions with village chairpersons and WMA village representatives, some of participant explained that some district officials covertly include their cattle among those of encroachers and also have farms inside the conservation area. These participants narrated that they consequently delay providing support for WMA operations, in order to protect their livestock and crops.

Quote 32: “*Our district government leaders are also the ones who influence the ongoing encroachments in our WMA. Others are the ones who own farms and cattle inside our WMA and they do it using the encroachers around by combining their cattle”* P1, FGD 2.

However, during interviews with the District Game Officers, they explained that elected officials at government local level, particularly village chairpersons, often spread rumour among community members about district officials, including the District Commissioner. They do this to make the community distrust the officials, so that when these officials emphasize the importance of conservation, the community dismisses their efforts, believing that the officials themselves are involved in the destruction of the WMA. These contrasting statements are difficult to reconcile creating considerable ambiguity:

Quote 33: *We were once given rumours that the District Commissioners and the Game Officers were pretending to protect, while they owned large farms inside the WMA and cattle. This even gained attention from the media, but all of this was aimed at [distracting from] and weakening conservation efforts.* IDI 8.

###### Lack of regular patrols and satisfactory equipment

Further, Village Game Scouts groups and WMA management committee members explained that there are limited numbers of patrols, which are inadequate following the influx of new immigrants who have encroached on the conservation area. An NNP stakeholder added that the WMA lacks sufficient equipment for operations to evict people from the conservation area including firearms and vehicles:

Quote 34: “*Our area is big, so it is hard for us to cover the whole area without transport. Previously we had no firearms, so it was really hard. We even got attacked by the livestock keepers because we only had batons for the patrols, so sometimes we were even scared to conduct patrols frequently”.* P2, FGD 6.

###### Lack of support from the Tanzania Forestry Services for operations

A stakeholder from KGR, a District Game Officer, and one Village Game Scout participant explained that a large portion of the charcoal used in Ifakara town comes from the WMA conservation area. They mentioned that some charcoal burners possess licenses issued by TFS (Tanzania Forestry Services), yet TFS does not closely monitor where the charcoal burners source their wood or the tree species they cut for charcoal production.

Quote 35: *Most of the charcoal at Ifakara comes from ILUMA WMA. Surprisingly, Tanzania Forestry Services are the one who issues charcoal licenses and collects taxes from the natural resource gate at Kivukoni, while they are not sure where all this charcoal is coming from. They must at least be aware of all this to reduce the charcoal burning from the WMA area”.* IDI 2.

###### Upgrade of nearby protected areas

Just over half of IDI participants and one FGD group explained that the upgrade of the nearby protected areas contributed to increased encroachment in the WMA. KGR and NNP personnel explained that when the Selous Game Reserve was under TAWA, they often assisted with patrols within the WMA because they have authority over the WMAs. However, now that it is a national park with different management policies, there is less direct action in neighbouring WMAs. Consequently, their contribution to joint operations is low:

Quote 36: “*After the Selous was upgraded to Nyerere National Park in 2019, this is when the WMA started to be greatly encroached by this Sukuma tribe. We used to help patrols and sometimes we used the WMA roads on our routes, so people were scared a lot when they saw us, but now NNP has no power like we at TAWA have over the management of the WMA.”* IDI 2.

Additionally, a District Game Officer also explained that recent upgrade of the nearby KGCA to the KGR had also contributed to an increased livestock herding within the WMA. During the time the KGCA was operational, extensive livestock herding occurred within its boundaries. However, after the upgrade, the policies and practices applied across game reserves no longer allowed such activities, prompting many livestock keepers to move to the WMA. He/she explained that the WMA regulations are not as strictly enforced as those of the game reserve, where livestock are more consistently confiscated and auctioned off. So, herders moved into the WMA where they know they can just pay fines to retrieve their livestock:

Quote 37: “*Recently the KGCA has upgraded to KGR. The former management practices allowed human activities like livestock herding but the latter doesn’t anymore, so because TAWA are stable financially, with manpower and equipment, they have managed to remove the livestock keepers from that area. They now have moved to the WMA because they are aware that we do not have that power, so they take us lightly”*. IDI 8.

#### 3.2.3 Changes in community attitudes

Various stakeholders, particularly those at village level, reported changes in community attitudes towards the WMA (Table 7). Village chairpersons explained that, during the establishment process, each village was informed and agreed to become a WMA member village and the community had positive attitudes. However, it was noted by a participant from one village that the community there strongly disagreed, complained that they were not properly involved and did not agree to be a WMA member village. This was also reported in a stakeholder meeting, where one of the attendees explained that his/her village had been opposed to the WMA since its establishment:

**Table 7:** Proportion of participants who mentioned changes in community attitudes, based on transcript review and coding from both IDI and FGDs.

|  | Coding frequency<br>(n [Proportion]) |
| --- | --- |
| 3.2.3 Changes in community attitudes | Participants<br>(n= 87) |
| 3.2.3.1 *Change from positive to negative* | 43 [49%] |
| 3.2.3.2 *Change from negative to positive* | 6 [7%] |
| 3.2.3.3 *No change* | 16 [18%] |

Quote 38 *“In my village, you can’t talk anything about ILUMA and expect people to listen. The community insist that they do not recognize this WMA. They are saying that only the village chairperson was taken to Mahenge* [location of district government office in one of the WMA district] *to sign the agreements, without them being directly involved. In every meeting we hold, the villagers insist I bring the* [village] *chairperson who was in power at the time WMA was established, for him to explain the truth. The* [village] *chairperson is no longer in power and when I try to reach him, he doesn’t show up as he is also busy with fishing in ILUMA.”* P1, FGD 8.

Most perceptions expressed in FGDs indicated changes of community attitudes from positive to negative, while fewer (∼19%) noted a recent change from negative to positive. Meanwhile, one-quarter reported no changes at all. IDIs less frequently related perspectives on community attitudes, presumably because most of participants were at district or regional level and might not be as aware of these issues. However, a few of them, particularly those with the District Game Officers, also reported changes similar to those noted in the FGDs, including both long-term changes from positive to negative and also more recent changes from negative to positive.

##### 3.2.3.1 Changes from positive to negative perceptions

Village chairpersons, Village Game Scouts and WMA village representatives from one of the WMA member villages in Ulanga district reported that their community wished to withdraw from the WMA:

Quote 39: “*Personally, from my perspective and that of my village, we do not want ILUMA. Even if someone goes and holds a meeting there, you will hear what we are saying. It’s not like we are just saying this from our minds, no, it’s what is in our villages.*” P3, FGD 7.

Quote 40*: “In my village, people talk a lot of negatives about this WMA. When you get chance, come and talk even with few members. You will hear them start asking for help to get their land back*.*”* P1, FGD 6.

Participants explained that community have changed to negative attitudes due to not realizing any benefits from the WMA, either for community development support or even for conservation improvement on their land:

Quote 41*: “Maybe I can say two things, because it seems like my colleagues here are afraid to say. Seriously, the community attitude is highly negative because the agreement has become different. There is no money. I mean there is no income for the villages. There are none of those services we were being told before that we could get from the WMA, like schools or health centres. But also, ILUMA has been somewhere that people are farming, herding livestock, burning charcoal. There seems to be no conservation there.”* P3, FGD 7.

Others participant explained that, due to the illegal land use by recent immigrants, the long-established local communities who set their land aside for conservation felt upset because it was being used by others illegally. They shared their perceptions that these immigrants benefited from their land rather than they themselves, which led them to change their attitudes to negative ones:

Quote 42*: “Resentment truly exists; why a pastoralist from Shinyanga region comes to settle there and ends up being more successful than us, and we are farming out there in unfertile sandy soil. So, you just find sometimes people say ILUMA means nothing to them because they are hurting in that way.*” P3, FGD 6.

Furthermore, one village chairperson explained that they wished to withdraw from the WMA because they perceived their land was used only to benefits a few individuals. They explained that people who had authority over the WMA at the district level, together with the former WMA management committee, had secretly allowed encroachers to graze livestock within the WMA area in exchange for money.

Quote 43*: “ILUMA has been like somewhere that a few people generate income, especially our leaders at district level and that of our WMA. That is why we and our community have changed attitudes. It has been somewhere only a few individuals are benefiting from; they build their houses and do other things but the villages have been left with nothing while their land is being used*.” P3, FGD 7.

On the hand, others explained that the increase in local population is also among the reasons for these changes. There is increased need for land for agriculture and settlement, as the majority of the community depends on farming for their primary livelihood.

Quote 44: “*What causes them to think that now we can take over ILUMA WMA is that ten years is a long time and youths have now grown up. They want to get married and need settlements and farms. The population is continuing to increase, which is why they now think that it’s better to get their land back.*” P4, FGD 8.

One Village Executive Officer explained that in their village, the community had previously collected sand from an area that was later included in the WMA. However, due to changes in management, they are no longer permitted to do so and so, which led them feel that their rights are being denied:

Quote 45: “*The community is complaining about being denied access to the sand they used to extract from the WMA area. They feel oppressed and believe it would be better to claim their land back, so they can continue their activities freely*.” P5, FGD 8.

It was perhaps unsurprisingly that this was reported from the same village that claimed not to have been involved during the WMA establishment; so this might have occurred because the community was not fully aware of the new regulations implemented after the WMA was established.

##### 3.2.3.2 Recent changes from Negative to positive attitudes

Some other stakeholders reported recent changes from negative to positive attitudes among community members. They attributed these changes to a small amount of income that the WMA villages began to realize between a year of 2022 and 2023, together with changes in WMA leadership that occurred in October 2022:

Quote 46: *“Two years ago, they were rejecting ILUMA and didn’t want it because they didn’t see any benefits from conservation. But after this period of two years, during which they received little income from the WMA, they have changed their minds. They no longer speak negatively about ILUMA, and they expect to continue benefiting from it.”* P3, FGD 9.

Quote 47: “*Most villages were unhappy with the way the former WMA management committee were managing their WMA. They were arguing that their land only benefited those management committees while they themselves are not. We conducted a new election last year, at least now you can hear them saying, yes, we still want to be a WMA village member now.* IDI 3.

##### 3.2.3.3 No changes

On the other hand, one village chairperson and a WMA village representative, both from Kilombero district, reported no changes had occurred in their community attitudes towards the WMA, and many still wished to remain a member village.

Quote 48: “*My village has no problem with ILUMA WMA since it was established and they still have hope that it will benefit them, though complaints about these encroachers who freely farm on their land do exist.* P3, FGD 10.

However, one other participant explained that other community members feel somewhat trapped because they are aware that withdrawing from the WMA would neither return their land nor provide further benefits from it:

Quote 49: “*When the villagers become serious about reclaiming their land, they face obstacles. They told us that if we withdraw, the land will remain under the WMA, so we are left in a hard situation. It’s as if we have been forced to say we do like ILUMA. But if we had the power to withdraw and reclaim our land, ILUMA could have died a long ago.*” P3, FGD 7.

#### 3.2.4 Revenue sources and utilization

Interviews with District Game Officers and discussions with WMA management committee members indicated that the main source of revenues is from fines charged for unauthorized activities inside the conservation area, particularly livestock herding (Table 8). The second most important income source was the collaborative agreement with researchers from IHI, SUA and UCC who collaborated with the WMA since 2021, followed by additional access fees paid by research interns added to the project. The least important income stream was the small number of regular tourist visitors. These participants also explained that ILUMA had not yet attracted an investor, other than one recently engaged party with whom they had signed a contract but who had not yet contributed any income to the WMA:

**Table 8:** Revenue sources from when the new WMA leadership held in power Oct 2022-Dec 2023.

| Revenue sources | Amount (Tsh/US\$) |
| --- | --- |
| Fines | Tsh 102,130,000 [\$39,281] |
| Collaborative agreement with researchers | Tsh 24,000,000 [\$9,231] |
| Additional access fees for research interns | Tsh 18,000,000 [\$6,923] |
| Tourists | Tsh 11,866,000 [\$4,564] |
| Total | 155,996,000 [\$59,998] |

Quote 50: “*We do not have many revenue sources. Most are fines, which I can say is illegal income because we get from mostly the livestock keepers. Other revenues are from researchers from Ifakara Health Institute, [and SUA and UCC] who are the ones who give us support for conducting patrols. We have no investors but only very few tourists and students visiting our area.”* P2, FGD 14.

Note, however, that at the time of these IDIs and FGDs, the WMA accountant reported that they were not satisfactorily informed of revenues collected in previous years, as they only have one year and a few months of records since they were appointed as WMA management committee, so they only provided a report on covering the period since their appointment to the WMA management committee in October 2022 (Table 8).

Only two participants, one village chairperson and an NNP representative, perceived the revenue use in the WMA as being reasonably distributed but simply inadequate in scale, relative to community needs and conservation operations running costs:

Quote 51*: “ILUMA doesn’t have much revenue and has contributed very little because it lacks investors for hunting activities. However, it doesn’t perform well in benefiting the community because the area itself still requires a lot of funds for its conservation.”* IDI 1.

Overall, participants demonstrated varied perspectives about the effectiveness of revenue usage (Table 9), with notable differences in perceptions based on leadership tenure. Some participants expressed no concerns about revenue distribution, while others linked effectiveness or ineffectiveness to specific leadership periods. These varied perspectives showed how ineffective use of revenue may lead to community disappointment and reduced motivation to support the WMA.

**Table 9:** Perspectives on revenue use effectiveness for conservation and community support functions.

| 3.2.4. Revenue sources and utilization patterns | IDI | FGD |
| --- | --- | --- |
|  | (n=9) | (n/N=78/16) |
| 3.2.4.1. Perceptions of effective revenue utilization |  |  |
| • Effective only after the formation of new leadership | 1 [11%] | 4 [25%] |
| • Effective | 1 [11%] | 1 [6%] |
| 3.2.4.2 Perceptions of ineffective revenue utilization |  |  |
| • Ineffective | 0 [0%] | 1 [6%] |
| • Ineffective specifically during the tenure of former WMA management committee | 1 [11%] | 5 [31%] |

##### 3.2.4.1 Perceptions of effective revenue utilization

Participants in one quarter of FGDs and one IDI expressed views that the new WMA management committee at the time were using their minimal revenue fairly and transparently. Even though the revenue shared with the community amounted to only two million Tanzanian shillings ($769) per village, this was noted by some as an important gesture that was six times greater than any payments received previously:

Quote 52*: “The way that they have started, it’s good and if they become more experienced with leadership skills, and avoid the greed of money, and ensure stakeholders involvement, ILUMA is going to change. We can now say something because they have reported the revenues and expenditures, we have observed something that even the village chairpersons have appreciated, and each village has received two million thus far [this year]”* IDI 3.

##### 3.2.4.2 Perceptions of ineffective revenue utilization

More FGDs than IDIs revealed perceptions that the use of WMA revenue for community development and conservation activities was ineffective, with nearly one-third expressing concerns. They particularly pointed out the former WMA management committee, who apparently remained in place for over ten years since the establishment of the WMA, without any fresh elections over that period:

Quote 53: “*In the past ten years under the former management committee of ILUMA WMA, of whom I cannot talk too much, revenue seemed not to be used effectively because, even when they left from power and handed over bank statements, the balance was zero, empty, meaning there was nothing. Now you ask yourself, what were they doing all those ten years? No benefits to the villages but only for the individuals!”.* IDI 3.

Similarly, this was noted at a stakeholders meeting, where a WMA financial report was presented. This lack of transparency and financial mismanagement contributed to the decision made at the meeting for the WMA leadership to be changed, which occurred one month later through an election overseen by the district authorities:

Quote 54: “*Truly money killed Jesus Christ. Looking at this financial report, expenditures were mostly used for allowances. This is not right. There is nothing the community has benefited from. Each statement here talks about allowances for either transport or meetings. What was it discussed in all of these meetings, with management committee only present and without other stakeholders?”* District representative at public stakeholder meeting.

#### 3.2.5 Stakeholders recommendations for the sustainability of the ILUMA WMA

The stakeholders involved in this study have put forward several key recommendations aimed at ensuring the long-term sustainability of the ILUMA WMA. These recommendations focused on enhancing stakeholder involvement, governance and financial management, the effectiveness of patrols, infrastructure development, conservation education programs, and developing marketing strategies. Additionally, they highlighted the need for government and donor support, capacity building for WMA management committee and community leaders, and exploring alternative income sources. By addressing these areas, stakeholders believe that the WMA can achieve its conservation goals and provide meaningful benefits to local communities (Table 10).

**Table 10:**
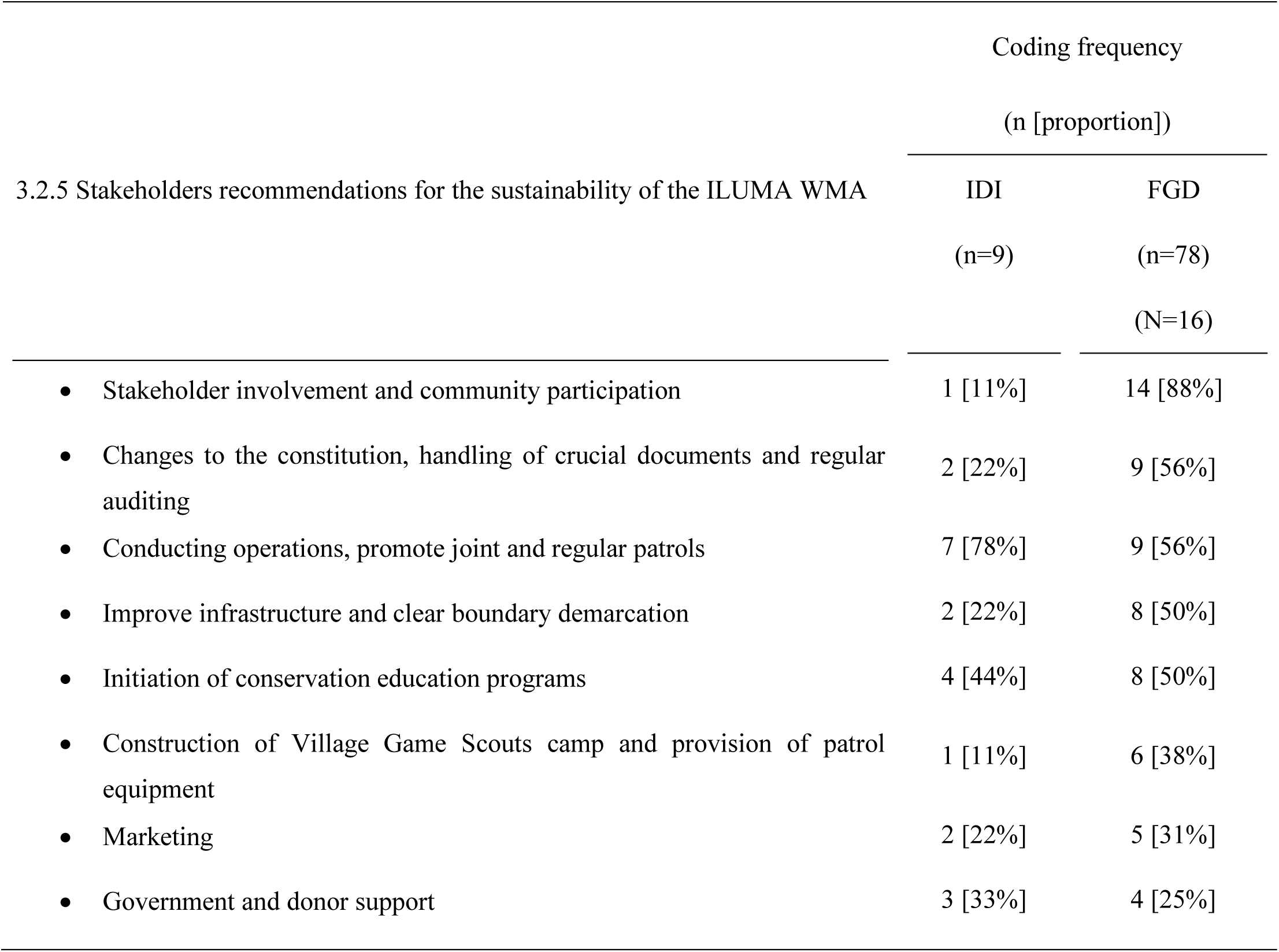
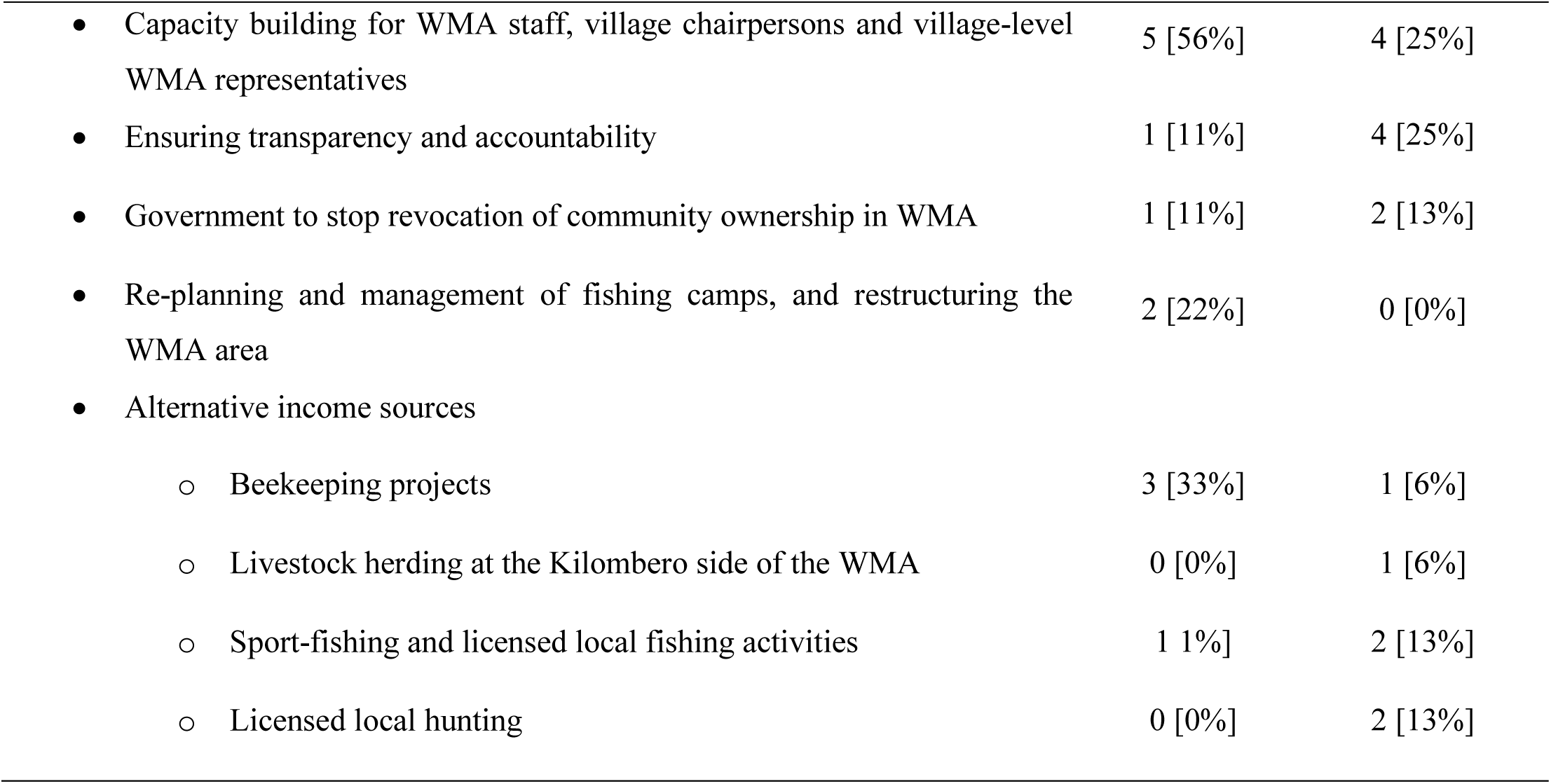
Stakeholders suggestion regarding achieving sustainability.

| 3.2.5 Stakeholders recommendations for the sustainability of the ILUMA WMA | Coding frequency |  |
| --- | --- | --- |
|  | (n [proportion]) |  |
|  | IDI<br>(n=9) | FGD<br>(n=78)<br>(N=16) |
| • Stakeholder involvement and community participation | 1 [11%] | 14 [88%] |
| • Changes to the constitution, handling of crucial documents and regular auditing | 2 [22%] | 9 [56%] |
| • Conducting operations, promote joint and regular patrols | 7 [78%] | 9 [56%] |
| • Improve infrastructure and clear boundary demarcation | 2 [22%] | 8 [50%] |
| • Initiation of conservation education programs | 4 [44%] | 8 [50%] |
| • Construction of Village Game Scouts camp and provision of patrol equipment | 1 [11%] | 6 [38%] |
| • Marketing | 2 [22%] | 5 [31%] |
| • Government and donor support | 3 [33%] | 4 [25%] |
| • Capacity building for WMA staff, village chairpersons and village-level WMA representatives | 5 [56%] | 4 [25%] |
| • Ensuring transparency and accountability | 1 [11%] | 4 [25%] |
| • Government to stop revocation of community ownership in WMA | 1 [11%] | 2 [13%] |
| • Re-planning and management of fishing camps, and restructuring the WMA area | 2 [22%] | 0 [0%] |
| • Alternative income sources |  |  |
| ○ Beekeeping projects | 3 [33%] | 1 [6%] |
| ○ Livestock herding at the Kilombero side of the WMA | 0 [0%] | 1 [6%] |
| ○ Sport-fishing and licensed local fishing activities | 1 [1%] | 2 [13%] |
| ○ Licensed local hunting | 0 [0%] | 2 [13%] |

##### Stakeholder involvement and community participation

Almost all FGD groups, both at the village and district levels, together with an IDI at the regional level, emphasized the crucial importance of fostering stronger participation, involvement, and relationships among key stakeholders, including the WMA management committee, district advisory board members, WMA village representatives and village chairpersons, to ensure improved performance of the WMA. One district advisory board member and one regional stakeholder voiced concerns regarding the current situation, expressing the perception that ILUMA operates as an independent institution solely managed by a small number of selected community members, namely the WMA management committee. Due to that, there is a lack of teamwork and collaboration with government stakeholders. Specifically, the village chairpersons tended to view the WMA as just another separate institution, while the WMA management committee see themselves as the primary decision-makers and owners of the WMA. They explained that, in their view, WMA activities are consequently being carried out without proper consultation or involvement of all relevant stakeholders, potentially leading to resource mismanagement and ineffective decision-making:

Quote 55*: “The village chairperson is among the elected community representatives, but today you find the chairperson appearing as an ordinary and ignorant person. Let me tell you something, let’s convey a message to them that we village chairpersons have the power to destroy significantly, but also we have the power to build immensely. If there will be no involvement, this WMA is going to die very soon, but if they acknowledge our importance as chairpersons, this WMA will bring fruits from generation to generation*”. P5, FGD 10.

In addition to involving stakeholders, one community leader recommended that there was a critical need to engage all local community members regarding WMA matters. He reported that many individuals within the community feel disconnected and have lost their sense of ownership over their land:

Quote 56: *“That sense of ownership is completely absent; the community don’t see if conserving the WMA is their responsibility. And as it’s known, the first guardians in conservation are the communities nearby, and not the rangers with the firearms and so on.”* P2, FGD 13.

##### Need for improved document management, constitutional changes and financial auditing

Discussions with the majority of community leader groups and of district advisory board members emphasized the need for revisions to the WMA constitution to reflect the current situation better. A District Forest Officer pointed out aspects, such as financial management, meeting schedules, and measures against illegal activities are required to be revised.

Quote 57: *The WMA themselves must make sure they review their constitution. Things have been changed, so now they have to go through the constitution and see what should be added or removed”* P1, FGD 16.

Additionally, almost all village chairpersons and district advisory board members reported that important documents like by-laws, and contracts related to the WMA were missing from their offices. These participants recommended that it’s crucial to ensure these documents are distributed and made easily accessible to all the stakeholders.

Quote 58*: “In our village offices we do not have any document related to ILUMA. How can we even know what is it about? We need to have all the required documents for us to be aware*.” P3, FGD 7.

The District Legal Officer emphasized the importance of conducting regular financial audits of this WMA. She pointed out that, according to WMA regulations, these audits are required but have not been properly carried out, potentially leading to issues with financial management and accountability within the WMA.

Quote 58*: I wonder how can we have an institution that collects money for the community while there is no auditing of those who manage those finances. We never heard about that until in the stakeholders meeting, where we heard the auditing should have been done. Meaning it wasn’t being done for the past ten years! Auditing is very important, and we have to know the feedback so that we can be sure that our finances are well-managed”.* P2, FGD 16.

##### WMA patrols, infrastructures, and demarcations

WMA management committee members, Village Game Scouts, WMA village representatives and district advisory board member all explained that major operations to remove encroachers who have settled within the WMA were urgently needed, and that these then needed to be followed by regular patrols. The District Game Officers explained that they have been looking for permission for such operations since the year 2022, but had found this challenging due to the need for central government involvement. Since these operations require the coordination of various personnel beyond the WMA, including police officers, TANAPA, TAWA and TFS, orders must come from the central government. They explained that it has been a complicated procedure as the process involves multiple levels of approval: the relevant districts must request permission from the regional government, which in turn must seek it from the national level, and the final permit must be signed either by either the president office or the ministry of natural resources.

Quote 59: *Since last year, we have been requesting the operation permit for evicting people from the conservation area. The problem lies within our government system, as there are a lot of complicated procedures to follow. We have submitted letters but we are still being told to wait until the Ministry of Natural Resources signs it, and we still haven’t received the permit”.* IDI 3.

Additionally, they explained that operations could be seen to have compromised human rights, so they need to be well-documented, widely communicated, and properly approved at the national level to ensure transparency and legality. They said that without proper approvals and public awareness, the WMA can face subsequent legal and ethical challenges.

To facilitate effective operations activities, WMA management committee and Village Game Scouts explained the need for adequate equipment, such as vehicles, more firearms and permanent camps for Village Game Scouts inside the WMA. Additionally, they suggested the need for infrastructure improvement within the WMA, including clear demarcations of boundaries using roads, instead of only relying on concrete beacons, which are sometimes destroyed by encroachers to create legal ambiguity (Figure 7).

**Figure 7:**
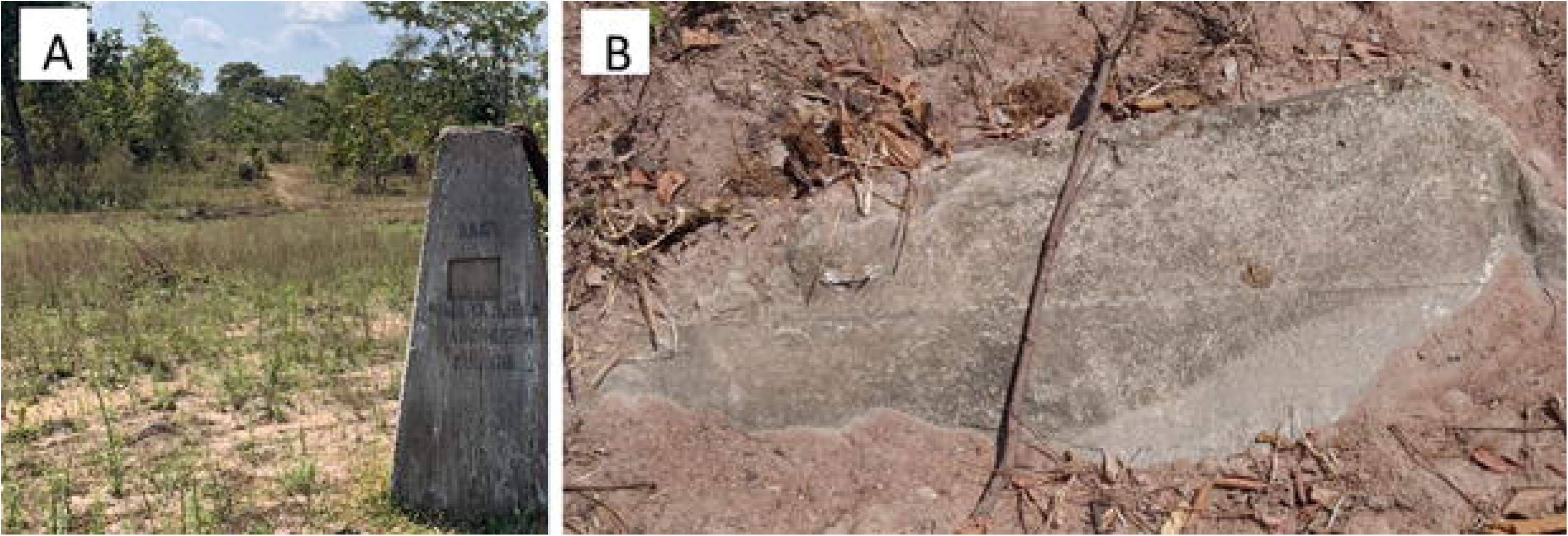
Images displaying the WMA beacon at the boundaries. Image A shows a standing beacon while image B shows an example of beacon that was buried by encroachers.

##### Conservation education programs and capacity building

Community leaders at villages and ward levels identified a significant gap in conservation education among the local community members. To address this, they strongly recommended initiating educational programs aimed at raising awareness about conservation, particularly targeting the younger generation.

These programs can play a role in illustrating the importance of conservation and empowering community members to actively participate in protecting their WMA. Furthermore, IDIs with KGR and NNP representatives emphasized the need to promote capacity-building for the WMA management committee, WMA village representatives and the village chairpersons, to help them become more effective in managing and protecting their WMA:

Quote 60*: Conservation also needs skills. I am not sure if the elected WMA representatives had received any training regarding conservation activities. They are just elected from among the village members, regardless of whether they are educated about conservation or not. Capacity building for them, and also the village chairpersons, is really important.”* IDI 1.

##### Marketing and alternative income sources

Participants mentioned initiatives such as beekeeping projects, setting up a livestock herding zone at the Kilombero side of the WMA, sport-fishing, licensed local fishing activities, and licensed local hunting as potential ways to diversify income for this WMA, rather than only depend on investors or donors:

Quote 61 *“For example, the WMA part in my village* [a village in Kilombero District] *could be used for grazing because I don’t think it has potential for other activities like tourism. We could use it for grazing to obtain permit fees for cattle herding, which could, in turn, help us finance other conservation activities and community development.”* P1, FGD 10.

##### Government support and stop from revocation of community ownership in WMA

Furthermore, some participants, including a regional government stakeholder, expressed the view that this WMA, which lacks investors and donor support cannot afford sufficient conservation operations, so they emphasized the need for better government support. This assistance should include resources and expertise from nearby protected areas such as NNP and KGR for patrol activities when necessary. Additionally, they emphasized the need for a policy review to ensure that nearby protected areas contribute funds to the WMAs and help them to meet their conservation goals, as these areas play a role in conserving wildlife outside their protected boundaries.

Quote 62*: “If you ask TANAPA to help with the WMA, they will tell you we have to follow the procedures! TAWA will say it’s not our role! I am saying this straightforward, the conservation institutions must*

*consider this. What was the aim of the WMA? It is because we failed to conserve wildlife that was outside our protected areas. The main aim was to conserve the wildlife and not directly to benefit the community. So how come now they make complicated processes in supporting the conservation of the WMA? They should have even set their funds aside for this struggling WMA to ensure its conservation.”* IDI 9.

Furthermore, some of district and regional stakeholders reported that there are unofficial rumours that this WMA might be upgraded to be a game reserve, or assimilated into NNP, so the community will lose its authority and rights regarding wildlife and other natural resources. One stakeholder strongly recommended that the WMA regulations should be reviewed, and that potential for punitive measures such as revocation of community ownership following conservation failures, should be reconsidered in favour of strategies aimed at capacity building and empowerment within local communities

Quote 63: “*I wonder why government thinks only about whether community fails in its conservation role? Their land will be taken instead of supporting their capacity to meet the goal. I think it is better to review the regulations, so that protected area near the WMA should take it on as part of their role, to ensure the community around meets the expectations from this WMA, because the land helps also to conserve the wildlife from such centrally managed protected areas like KGR and NNP.* IDI 8.

##### Re-planning and management of fishing camps, and restructuring the WMA area

Moreover, NNP and KGR stakeholders recommended restructuring the WMA, with a focus on returning highly encroached areas to villages for alternative uses rather than conservation purposes. They argued that it’s more practical to have a smaller WMA area that can effectively ensure conservation, rather than a larger one with limited conservation capacity. Additionally, one of them argued that the WMA implements no proper management of the fishing activities in their area, so they must ensure the authorized fishing camps (Duggan et al., 2024b) are monitored to reduce illegal fishing for the good sustainability.

Quote 64: “*The situation is not good, those large farms where trees have been cut down, people have already started living there, meaning that we will experience conflicts, and it’s very difficult to consider it as a conserved area now due to its capacity. It would be more appropriate if they will redesign, and those heavily encroached areas to be removed from the WMA because it has been like a grandmother farm. Some areas people buy beers inside there, like seriously! Where is conservation then?”* IDI 1.

## 4. Discussion

This study of stakeholder perspectives on the functions of the ILUMA WMA identified key themes related to livelihood and community development functions, environmental and wildlife conservation functions, changing community attitudes towards the WMA, revenue sources, and ideas for improving the sustainability of the WMA. Generally speaking, the findings reported herein are compatible with evidence from WMAs elsewhere in Tanzania.

The finding that employment opportunities for a small number of individuals appeared to be the only form of direct income accrued by community members echoes those of Funk (2015) and Kicheleri et al., (2018) from Burunge WMA. Overall, the community stakeholders who formed the ILUMA WMA appear to have realized negligible benefit at household or community levels since its establishment 13 years ago. This contrasts with findings regarding two other WMAs in Tanzania, namely Burunge and Ikona, where well-developed tourism activities supported community projects like building of schools, village offices and roads (Kicheleri et al., 2018; Kimario et al., 2020; Mgonja, 2023). However, progression from establishment through to improved household income and poverty reduction remains elusive for many WMAs (Bluwstein et al., 2018; Keane et al., 2020; Mgonja, 2023; Msangeni et al., 2024; Mwakaje, 2008; Noe & Kangalawe, 2015; Pailler et al., 2015), with benefits typically remaining insufficient to offset the wildlife induced costs accrued by the local communities (Lwankomezi et al., 2021).

A study of eight operational WMAs across Tanzania found that those with NGO, investor or donor support were more successful in supporting community projects, while those without such support struggled (Kimario et al., 2020). This shows that the success of WMAs in delivering benefits to local communities is heavily reliant on external support and investment to establish sustainable income streams. A study of one WMA (Mariki, 2019) indicated that cessation of donor and investment activities led to a failure to benefit local stakeholder communities. Therefore, beyond financial dividends and community project support, there is a need of rethink how local communities can benefit from direct access of natural resources such firewood, building materials, grasses, game meat on a sustainable basis.

Although the ILUMA WMA was perceived by some to have made positive contributions to conservation, it was generally perceived to have been only partially successful, with many noted failures. The apparent initial reduction in unauthorized activities following the ILUMA WMA establishment in 2011 that was narrated by participants aligns well with encouraging early experiences in Randilen WMA that were attributed to management changes in land use practice (Lee & Bond, 2018). However, consistent with recent ecological surveys (Duggan, 2023; Duggan et al., 2024a), participants in this study also explained that there was an increase in human encroachment from 2018 onwards. The village land that was combined to form this WMA was historically rich in wildlife, with intact forest (Brehony, 2005) but participants noted that the conservation status of the area was now worse than before the WMA was established. Interestingly, these unauthorised human activities have not only negatively affected the conservation status of the area but also traditional cultural practices within the forests which have now disappeared in some places. This emphasizes the need for conservation initiatives to consider the preservation of cultural sites and practices along with ecological goals.

Findings show that the community member in this area are generally not very well engaged in conservation efforts because they have not realized the benefits they expected through the legal mechanisms of the WMA, so some of them have resorted to illegal alternatives. For example, the narrative that village chairpersons had unofficially allocated portions of WMA land to new immigrants in exchange for illicit payments is consistent with historical experiences in the same area from before ILUMA was established, some of which culminated in the murder of village chairpersons (Brehony, 2005). These experiences in the ILUMA WMA contrast with the findings of another study in the MBOMIPA WMA, which evidenced a positive contribution of the WMA to biodiversity conservation attributed to the direct involvement of community members in reporting illegal activities because of benefits they accrued from the WMA (Mdete, 2016; Nebbo, 2015). Interestingly, similar observations have been made regarding the authorized fishing villages along the banks of the Kilombero River in ILUMA, where improved environmental and wildlife conservation was clearly documented and attributed to the custodian role played by these carefully regulated settlements (Duggan et al., 2024b). In this focal area of ILUMA, where these small resident communities were motivated by conditional access to a productive fishery, it appears that when such local communities are allowed to really benefit from the natural ecosystem, they are happy to help conserve it.

There are relatively few studies assessing the conservation achievements of WMAs, and other authors have emphasized the need for more critical assessment of this promising model for devolved conservation (Lwankomezi et al., 2021; Lee, 2018; Lee & Bond, 2018). Nevertheless, a comprehensive study of 13 operational WMAs established between 2003 and 2008 confirmed that WMAs may not be as effective as hoped in achieving their conservation goals (Dancer, 2013), as they still lack direct decision-making power for many of their operations (Bluwstein et al., 2016; Wright, 2017), just as we have reported here for ILUMA. We found that lack of sufficient financial resources for covering the costs of patrols a8nd operations was considered to be the most important reason why ILUMA has not yet achieved its conservation goals. ILUMA lacked adequate income and had no investors or donor support beyond the initial short-term establishment phase over the eight years since it was granted user rights. Similarly, a study of the *Wami Mbiki* WMA demonstrated that it also struggled to cover operational costs for conservation and consequently experienced increased encroachment after donor support for WMA activities ended (Mariki, 2019). Additionally, it was noted from the discussions with one regional-level participant this study that the *Wami Mbiki* was taken out of community ownership and converted into a game reserve by the government in 2021, specifically because it had failed to meet its conservation goals as a WMA.

Consistent with several studies (Butt, 2014; Green et al., 2019; Lesorogol & Lesorogol, 2024), more than half of IDIs in this study indicated that the protection efforts in nearby centrally managed protected areas, particularly the new KGR, increased pressure on the WMA, especially through livestock encroachment. The ILUMA WMA struggled to cope with this additional pressure, due to its limited finances and equipment, adding further to the evidence that WMAs are often vulnerable to encroachment pressures displaced from such centralized protected areas (Mariki, 2019; Keane et al., 2008; Rija, 2017). Given the role of WMAs as buffer zones for such parks and reserves, there seems to be a clear need for overhauling policy and practice to improve cooperation between these complementary conservation authorities.

Various stakeholders particularly those at village level reported changes in community attitudes towards the ILUMA WMA. Indeed, some village chairpersons expressed a wish for their village to step out from WMA membership, mostly due to unmet WMA objectives, lack of revenue and, to a lesser extent, financial mismanagement. These findings support the view that communities are less likely to support conservation when they do not see direct, tangible benefits that offset the conservation costs imposed on them including restricted access to valuable natural resources (Mariki, 2019; Hernold, 2020; Kimario et al., 2020; Mgonja & Uswege, 2022).

A further important recurring narrative was the perception that village land was now of greater *de facto* benefit to immigrants from elsewhere in Tanzania, who have encroached upon the area for livestock grazing, farming and even established permanent settlements. In the current era of growing rural migration (Salerno et al., 2024), this highlights the growing importance of understanding the fragmentation of communities and nuanced politics of identity and ownership within a given locale.

Additionally, the lack of communication and participation among relevant stakeholders remain a limitation for most WMAs in Tanzania (Kimario et al., 2020), a situation also reported herein for the in ILUMA WMA. Improving communication, to provide a clear consistent, up-to-date information about the WMA to the community members, and whereby addressing perceptions of inequity or unfairness are essential to rebuild trust and local support for this devolved access fees for ongoing conservation initiatives.

Findings that the ILUMA relies mainly on fines and access fee for ongoing research activities as its main sources of income is consistent with a national-scale evaluation of WMA performance (CWMAC, 2019). A complementary study has shown that WMAs need substantial initial capital investment from the relevant district government and government conservation agencies, to support WMA activities over the short-to-medium term, while also building capacity and expanding their income sources to achieve long-term sustainability (Kaswamila, 2012; Wilfred, 2010). Reports on financial viability of Tanzanian WMAs indicate that, for better developed WMAs like Burunge, revenues come mainly from tourism activities, although this exceptional success story could be attributed to partly due to its accessible location within the northern tourism circuit (Tang’are & Mwanyoka, 2023; USAID, 2016). Despite the generally discouraging picture for most other WMAs thus far, several studies emphasize the need to explore strategies for diversifying income sources, to ensure their long-term financial sustainability (ESPA, 2017; Msangeni et al., 2024; Nebbo, 2015; USAID, 2013). For instance, in areas with miombo woodland and forest resources like ILUMA could develop activities such as sustainable timber, sustainable charcoal, beekeeping (USAID, 2016) and, perhaps most promising of all, international carbon finance [https://carbontanzania.com/].

Our findings that a new management committee, chosen through election that was held soon after the first public stakeholder meeting in 2022, achieved notable improvements in how revenues were utilized emphasizes the importance for WMAs to adhere to their constitutional guidelines for governance and management. Also, the perceived need for relevant district authorities to implement regular financial audits, to ensure open accountability, is consistent with findings from other studies, showing that even WMAs with robust income streams, such as Ikona and Burunge, face challenges related to financial transparency and benefit-sharing conflicts (Kimario et al., 2020; Kisingo & Kideghesho, 2020; Nebbo, 2015; USAID, 2013). Encouragingly, even the tiny recent revenue shares from the ILUMA WMA allocated to member villages was considered by several participants to represent an important gesture and proof of principle.

Of course, this study had substantive limitations that should be considered when assessing the certainty and representativeness of the evidence generated. Although every effort was made to solicit perspectives from a broad spectrum of ILUMA WMA stakeholders at national, regional, district and village levels, based on ethnically neutral *a priori* criteria, the de facto outcome of this participant selection process was that no voices from any of the relatively new pastoralist communities in the area were included. Future research should prioritize the inclusion of this community to capture their distinct insights and experiences, to ensure more inclusive and representative findings. Furthermore, researcher bias is a common issue in many scientific fields, often arising from personal interests in reaching particular study conclusions (Ioannidis, 2005; Sarewitz, 2012). Qualitative social studies are especially susceptible to this bias, and many do not meet the most rigorous criteria established for sociological research (Paulhus, 1991; Schumm, 2021; Tong et al., 2007). Since data collection for this study was conducted by the first author, who had prior knowledge of the challenges faced by the WMA, there is therefore a possibility that investigator bias may have unintentionally been communicated to the participants. Additionally, due to time constraints, the investigator was unable to return the transcripts and findings to the participants for their review and feedback, which could have helped ensure the accuracy and validity of the data.

## 5. Conclusions

Despite these study limitations, this study nevertheless clearly identifies several challenges that the ILUMA WMA faces as it struggles to achieve its dual objectives of advancing conservation while also delivering benefits to stakeholder communities, the most important of which appears to be securing sufficient income. Despite the small income that this WMA has generated thus far, it has nevertheless managed to support conservation in the area and provide some token support for community development activities. This WMA now needs to urgently diversify and expand its income streams, so that it can cover its full operating costs and generate enough of a surplus to benefit local communities.

Interestingly, a recent study has shown that regulated authorized fishing settlements within the area, which are allowed direct but regulated access to natural resources, contribute positively to wildlife and forest conservation (Duggan et al., 2024b). Therefore, ILUMA should consider developing similar practices that directly involve the community members in such resident custodian roles that allow them conditional, carefully managed access to natural resources, such as seasonal grazing (Duggan, 2023; Duggan et al., 2024b). Livestock grazing has been identified as a key challenge for WMAs and other protected areas across Tanzania (Kimario et al., 2020), and recent evidence suggests that it could be compatible with wildlife conservation if astutely practiced and carefully regulated (Liang et al., 2018; Mtimbanjayo & Sangeda, 2018; Odadi et al., 2011; Odadi et al., 2011a; Tyrrell et al., 2020; Xu & Butt, 2024). Therefore, it is important to explore positive ways to integrate pastoralists and agro-pastoralists into conservation efforts by addressing their needs while also enforcing key conservation regulations, perhaps with stronger deterrents like outright confiscation. For instance, rainy season access to grazing within the WMA, at the time when pastoralists most urgently need to move their cattle off agricultural land and they can be most easily accommodated outside of the main tourist season (Duggan, 2023; Duggan et al., 2024b), might represent a more easily regulated, mutually beneficial compromise between these relatively recently arrived communities and the long-established villages who own and run the ILUMA WMA than the clear conflicts over land currently ongoing.

## Supporting information

Appendix 1 discussion guide tool

Appendix 2 participant consent form

## AUTHOR CONTRIBUTIONS

Lucia J. Tarimo: Conceptualization, investigation, methodology, implementation, formal analysis, writing original draft of the manuscript. Deogratius R. Kavishe: Methodology, review, editing, validation. Fidelma Butler: Conceptualization, review, editing, validation. Gerry F. Killeen: Acquisition of fund, conceptualization, methodology, implementation, review, editing and validation. Felister Mombo: Conceptualization, investigation, methodology, implementation, review, editing and validation. All authors read and approved the final submitted version of the manuscript.

## ACKNOWLEDGEMENTS

The authors wish to thank all the stakeholders of the ILUMA WMA for their invaluable participation in this study. A very special word of thanks is due to our recently deceased friend and colleague, Mr Octavian Malopola, without whom this work would never have even begun. We also thank Dr. Emmanuel Kaindoa, Mr. Frederic Masanja, and Mr. Fadhili Songo for the essential institutional support provided by the Ifakara Health Institute throughout the course of the study.

## SUPPORTING FILES

**Supplementary file 1;** Appendix 1: Discussion guide tool used for both focus group discussions and the in depth interviews https://doi.org/10.5281/zenodo.14063184

**Supplementary file 2**; Appendix 2: Participant consent form provided to the participant before the discussions https://doi.org/10.5281/zenodo.14063329

## REFERENCES

Agrawal, A., & Gibson, C. C. (1999). Enchantment and disenchantment: The role of community in natural resource conservation. World Development, 27(4), 629–649.

Lwankomezi, E., Kisoza, J., & Patrobas Mhache, E. (2021). Benefit Sharing in Community Based Conservation Programs: The Case of Makao Wildlife Management Area, Tanzania. EAST AFRICAN JOURNAL OF EDUCATION AND SOCIAL SCIENCES, Issue 2 (April to June 2021), 41–50. 10.46606/eajess2021v02i02.0074

Mariki, S. (2019). Successes, Threats, and Factors Influencing the Performance of a Community-Based Wildlife Management Approach: The Case of Wami Mbiki WMA, Tanzania. In J. R. Kideghesho & A. A. Rija (Eds.), Wildlife Management—Failures, Successes and Prospects. IntechOpen. 10.5772/intechopen.79183

Baldus, R. D., & Cauldwell, A. E. (2004). Tourist hunting and it’s role in development of wildlife management areas in Tanzania. Dar Es Salam. http://wildlife-baldus.com/download/Tourist%20Hunting%20in%20TZ%20-%20PART%20I.pdf

Balint, P. J. (2006). Improving community-based conservation near protected areas: The importance of development variables. Environmental Management, 38, 137–148.

Bluwstein, J., Homewood, K., Lund, J. F., Nielsen, M. R., Burgess, N., Msuha, M., Olila, J., Sankeni, S. S., Millia, S. K., & Laizer, H. (2018). A quasi-experimental study of impacts of Tanzania’s wildlife management areas on rural livelihoods and wealth. Scientific Data, 5(1). https://www.nature.com/articles/sdata201887

Bluwstein, J., Moyo, F., & Kicheleri, R. P. (2016). Austere conservation: Understanding conflicts over resource governance in Tanzanian wildlife management areas. Conservation and Society, 14(3), 218–231.

Brehony, Dr. E. (2005). A Study On Conflict In Ulanga District ,Morogoro Region Tanzania (p. 66).

Brockington, D. (2002). Fortress conservation: The preservation of the Mkomazi Game Reserve, Tanzania. Indiana University Press.

Butt, B. (2014). The political ecology of ‘incursions’: Livestock, protected areas and socioecological dynamics in the mara region of Kenya. Africa, 84(4), 614–637.

Byrne, D. (2022). A worked example of Braun and Clarke’s approach to reflexive thematic analysis. Quality & Quantity, 56(3), 1391–1412.

Chisanga, A. (2016). What explains success and failure in Community Based Natural Resource Management? A comparison of Botswana and Zambia [Master’s Thesis]. https://gupea.ub.gu.se/handle/2077/44849

Clarke, V., & Braun, V. (2017). Thematic analysis. The Journal of Positive Psychology, 12(3), 297–298.

CWMAC. (2019). Wildlife Management Area (WMA) Performance Assessment Report. https://pdf.usaid.gov/pdf_docs/PA00X5TW.pdf

Dancer, A. (2013). Do Community-conserved Areas in Tanzania Achieve Conservation Goals?: An Initiative-wide Study Using Remote Imagery and Matching Methods [PhD Thesis, Citeseer]. https://citeseerx.ist.psu.edu/document?repid=rep1&type=pdf&doi=90f1e0df18bbb9ec5afd2e1b878602ccfc98e660

Dressler, W., Büscher, B., Schoon, M., Brockington, D. A. N., Hayes, T., Kull, C. A., McCarthy, J., & Shrestha, K. (2010). From hope to crisis and back again? A critical history of the global CBNRM narrative. Environmental Conservation, 37(1), 5–15.

Duggan, L. (2023). The influence of community-defined land use plans and de facto land use practices on the relative abundance and distribution of large wild mammals in a community-based Wildlife Management Area in Southern Tanzania. University College Cork.

Duggan, L. M., Tarimo, L. J., Walsh, K. A., Kavishe, D. R., Crego, R. D., Elisa, M., Mombo, F., Butler, F., & Killeen, G. (2024b). An effective model for community-based conservation around authorized fishing settlements inside a devolved Wildlife Management Area in southern Tanzania. bioRxiv, 2024–07.

Duggan, L. M., Tarimo, L. J., Walsh, K. A., Kavishe, D. R., Crego, R. D., Elisa, M., Mombo, F., Butler, F., & Killeen, G. F. (2024). Direct comparative assessment of radial and transect surveys to document wild mammal activity across diverse habitat types. African Journal of Ecology, 62(3), e13309. 10.1111/aje.13309

Duggan, L., Walsh, K., Tarimo, L., Kavishe, D., Crego, R., Eliza, M., Butler, F., Mombo, F., & Killeen, G. (2024a). A subjective and intuitive approach to rapid, holistic assessment of natural ecosystem integrity across a community-managed conservation area in southern Tanzania. Authorea Preprints. https://essopenarchive.org/doi/full/10.22541/au.171733407.76333984

ESPA. (2017). Realising the promise of Tanzania’s Wildlife Management Areas (p. 4). https://www.espa.ac.uk/files/espa/Realising%20the%20promise%20of%20Tanzania%20Wildlife%20Management%20Areas.pdf

Fortmann, L., Roe, E., & Van Eeten, M. (2001). At the threshold between governance and management: Community-based natural resource management in Southern Africa. Public Administration and Development, 21(2), 171–185. 10.1002/pad.156

Funk, S. (2015). Challenging the Win-Win Proposition of Community-Based Wildlife Management in Tanzania A case of Burunge WMA. University College London, UK. https://www.ucl.ac.uk/pima/docs/theses/Funk_MSc.pdf

Garner, K.-A. (2012). CBNRM in Botswana: The failure of CBNRM for the indigenous San, the village of Xai Xai and the wildlife of Botswana [PhD Thesis, University of Guelph]. https://atrium.lib.uoguelph.ca/handle/10214/4053

Gibson, C. C., & Marks, S. A. (1995). Transforming rural hunters into conservationists: An assessment of community-based wildlife management programs in Africa. World Development, 23(6), 941–957.

Goldman, M. (2003). Partitioned Nature, Privileged Knowledge: Community-based Conservation in Tanzania. Development and Change, 34(5), 833–862. 10.1111/j.1467-7660.2003.00331.x

Green, D. S., Zipkin, E. F., Incorvaia, D. C., & Holekamp, K. E. (2019). Long-term ecological changes influence herbivore diversity and abundance inside a protected area in the Mara-Serengeti ecosystem. Global Ecology and Conservation, 20, e00697.

Hernold, H. (2020). Burunge Wildlife Management Area and effects on the villages around-: A case study in Babati district, Tanzania.

Hulme, D., & Murphree, M. (2001). African wildlife and livelihoods: The promise and performance of community conservation. https://www.cabidigitallibrary.org/doi/full/10.5555/20013145442

Ioannidis, J. P. (2005). Why most published research findings are false. PLoS Medicine, 2(8), e124.

Jones, B. (2010). The evolution of Namibia’s communal conservancies. *Community Rights*, Conservation and Contested Land, 119–133.

Kajembe, G. C., Monela, G. C., & Mvena, Z. S. (2006). Making community-based forest management work: A case study from Duru-Haitemba village forest reserve, Babati, Arusha, the United Republic of Tanzania. http://www.treesforlife.info/fao/Docs/P/y4807b/Y4807B15.pdf

Kaswamila, A. (2012). An Analysis of the Contribution of Community Wildlife Management Areas on Livelihood in Tanzania. In A. Kaswamila (Ed.), Sustainable Natural Resources Management. InTech. 10.5772/32987

Kavishe, D. R., Msoffe, R. V., Malika, G. Z., Walsh, K. A., Duggan, L. M., Tarimo, L. J., Butler, F., Kaindoa, E. W. W., Ngowo, H. S., & Killeen, G. (2024a). A self-cooling self-humidifying mosquito carrier backpack for transporting live adult mosquitoes on foot over long distances under challenging field conditions. bioRxiv, 2024–04.

Kavishe, D. R., Walsh, K. A., Msoffe, R. V., Duggan, L. M., Tarimo, L. J., Butler, F., Govella, N. J., Kaindoa, E. W., & Killeen, G. (2024b). Comparative attractiveness of Anopheles quadriannulatus and Anopheles arabiensis to humans estimated by comparing the relative abundance of these two species in larval samples, unbaited adult catches and human-baited adult catches. bioRxiv, 2024–08.

Keane, A., Jones, J. P. G., Edwards-Jones, G., & Milner-Gulland, E. J. (2008). The sleeping policeman: Understanding issues of enforcement and compliance in conservation. Animal Conservation, 11(2), 75–82. 10.1111/j.1469-1795.2008.00170.x

Keane, A., Lund, J. F., Bluwstein, J., Burgess, N. D., Nielsen, M. R., & Homewood, K. (2020). Impact of Tanzania’s Wildlife Management Areas on household wealth. Nature Sustainability, 3(3), 226–233.

Kicheleri, R. P., Treue, T., Nielsen, M. R., Kajembe, G. C., & Mombo, F. M. (2018). Institutional Rhetoric Versus Local Reality: A Case Study of Burunge Wildlife Management Area, Tanzania. SAGE Open, 8(2), 215824401877438. 10.1177/2158244018774382

KILORWEMP PIU. (2016). Annual result report (p. 87) [Project report]. http://www.diplomatie.be/oda/23131_ENABEL_ANN_REPORT_TAN1102711_17_AnnualReport_2017-02-28_000_20170301100258.pdf

Kimario, F. F., Botha, N., Kisingo, A., & Job, H. (2020). Theory and practice of conservancies: Evidence from wildlife management areas in Tanzania. Erdkunde, 117–143. 10.3112/erdkunde.2020.02.03

Kisingo, A. W., & Kideghesho, J. R. (2020). Community Governance of Wildlife Resources: Implications for Conservation, Livelihood, and Improvement in Democratic Space. In J. O. Durrant, E. H. Martin, K. Melubo, R. R. Jensen, L. A. Hadfield, P. J. Hardin, & L. Weisler (Eds.), Protected Areas in Northern Tanzania (Vol. 22, pp. 113–120). Springer International Publishing. 10.1007/978-3-030-43302-4_8

Kiss, A. (2004). Is community-based ecotourism a good use of biodiversity conservation funds? Trends in Ecology & Evolution, 19(5), 232–237.

Kull, C. A. (2002). Empowering Pyromaniacs in Madagascar: Ideology and Legitimacy in Community-Based Natural Resource Management. Development and Change, 33(1), 57– 78. 10.1111/1467-7660.00240

Lane, M. B. (2001). Affirming New Directions in Planning Theory: Comanagement of Protected Areas. Society & Natural Resources, 14(8), 657–671. 10.1080/08941920118212

Larson, L. E. (1976). A History of the Mahenge (Ulanga) District [PhD Thesis, PhD thesis, University of Dar es Salaam]. https://www.researchgate.net/profile/Lorne-Larson/publication/34887155_A_history_of_the_Mahenge_Ulanga_District_ca_1860-1957/links/5804cdc508aee314f68e04d0/A-history-of-the-Mahenge-Ulanga-District-ca-1860-1957.pdf

Lee, D. E. (2018). Evaluating conservation effectiveness in a Tanzanian community wildlife management area. The Journal of Wildlife Management, 82(8), 1767–1774. 10.1002/jwmg.21549

Lee, D. E., & Bond, M. L. (2018). Quantifying the ecological success of a community-based wildlife conservation area in Tanzania. Journal of Mammalogy, 99(2), 459–464. 10.1093/jmammal/gyy014

Lesorogol, C., & Lesorogol, P. (2024). Community-Based Wildlife Conservation on Pastoral Lands in Kenya: A New Logic of Production with Implications for the Future of Pastoralism. Human Ecology, 52(1), 15–29. 10.1007/s10745-024-00482-9

Liang, W., Linxiu, Z., Min, W., Erustus, K., & Cong, D. (2018). The development of wildlife community conservancies in Kenya: A preliminary review. Journal of Resources and Ecology, 9(3), 250–256.

Mariki, S. B. (2018). Successes, threats, and factors influencing the performance of a community-based wildlife management approach: The case of Wami Mbiki WMA, Tanzania. In Wildlife Management-Failures, Successes and Prospects. IntechOpen London, UK.

Martin, R. B. (1986). Communal areas management programme for indigenous resources. WCED Archive Collection; v. 32, Doc. 235. https://idl-bnc-idrc.dspacedirect.org/bitstream/10625/2581/1/WCED_v32_doc235.pdf

Mbaiwa, J. E. (2004). THE SUCCESS AND SUSTAINABILITY OF COMMUNITY-BASED NATURAL RESOURCE MANAGEMENT IN THE OKAVANGO DELTA, BOTSWANA. South African Geographical Journal, 86(1), 44–53. 10.1080/03736245.2004.9713807

Mgonja, J. T. (2023). Assessing community perceptions about the contributions and impacts of Wildlife tourism to rural livelihoods: Wildlife management areas perspective. Tanzania Journal of Forestry and Nature Conservation, 92(1), 64–81.

Mgonja, J. T., & Uswege, D. N. (2022). Assessment of factors moderating community attitudes towards wildlife tourism and conservation: A case of Ikona and Makao wildlife management areas. Tanzania Journal of Forestry and Nature Conservation, 91(2), 214– 233.

Ministry of Natural Resources and Tourism. (2011). The development and implementation of an integrated management plan of Kilombero valley food plain Ramsar site (p. 63). http://www.diplomatie.be/oda/38221_ENABEL_FIN_REPORT_TAN0401111_24_RapFin_--_000.pdf

Msangeni, S., F Kess, J., & Mariki, S. (2024). Contribution of Wildlife-based Tourism to Household Income and Income Inequality: A Case of Burunge Wildlife Management Area in Tanzania. EAST AFRICAN JOURNAL OF EDUCATION AND SOCIAL SCIENCES, 5(1), 44–54. 10.46606/eajess2024v05i01.0348

Mtimbanjayo, J. R., & Sangeda, A. Z. (2018). Ecological effects of cattle grazing on Miombo tree species regeneration and diversity in Central-Eastern Tanzania. Journal of Environmental Research, 2(1), 1–7.

Murphree, M. (2004). COMMUNAL APPROACHES TO NATURAL RESOURCE MANAGEMENT IN AFRICA: FROM WHENCE AND TO WHERE? Journal of International Wildlife Law & Policy, 7(3–4), 203–216. 10.1080/13880290490883250

Mwakaje, A. G. (2008). Wildlife Management Areas in Tanzania: A Study of Opportunities and Challenges. Tanzania Journal of Development Studies, 8(2). https://www.ajol.info/index.php/tjds/article/view/60427

Nebbo, J. (2015). Wildlife management area strategy in sustainable conservation of wildlife resources, poverty reduction and in the mitigation of human/wildlife conflicts: The case of MBOMIPA in Iringa, Tanzania. [PhD Thesis, University of Eldoret]. http://41.89.164.27/handle/123456789/931

Nelson, F., & Agrawal, A. (2008). Patronage or Participation? Community-based Natural Resource Management Reform in Sub-Saharan Africa. Development and Change, 39(4), 557–585. 10.1111/j.1467-7660.2008.00496.x

Noe, C., & Kangalawe, R. Y. (2015). Wildlife protection, community participation in conservation, and (dis) empowerment in southern Tanzania. Conservation and Society, 13(3), 244–253.

Odadi, W. O., Jain, M., Van Wieren, S. E., Prins, H. H., & Rubenstein, D. I. (2011). Facilitation between bovids and equids on an African savanna. Evolutionary Ecology Research, 13(3), 237–252.

Odadi, W. O., Karachi, M. K., Abdulrazak, S. A., & Young, T. P. (2011). African Wild Ungulates Compete with or Facilitate Cattle Depending on Season. Science, 333(6050), 1753–1755. 10.1126/science.1208468

Pailler, S., Naidoo, R., Burgess, N. D., Freeman, O. E., & Fisher, B. (2015). Impacts of community-based natural resource management on wealth, food security and child health in Tanzania. PloS One, 10(7), e0133252.

Paulhus, D. L. (1991). Measurement and control of response bias. Measures of Personality and Social Psychological Attitudes/Academic Press, Inc.

Rija, A. A. (2017). Spatial pattern of illegal activities and the impact on wildlife populations in protected areas in the Serengeti ecosystem [PhD Thesis, University of York]. https://etheses.whiterose.ac.uk/20276/

Roe, D. (2011). Community-based natural resource management: An overview and definitions. CITES and CBNRM, 18.

Salerno, J., Gaughan, A. E., Warrier, R., Boone, R., Stevens, F. R., Keys, P. W., Mangewa, L. J., Mombo, F. M., de Sherbinin, A., & Hartter, J. (2024). Rural migration under climate and land systems change. Nature Sustainability, 1–10.

Sarewitz, D. (2012). Beware the creeping cracks of bias. Nature, 485(7397), 149–149.

Schumm, W. R. (2021). Confirmation bias and methodology in social science: An editorial. Marriage & Family Review, 57(4), 285–293. 10.1080/01494929.2021.1872859

Shackelton, S., & Campbell, B. (2000). Empowering communities to manage natural resources: Case studies from Southern Africa. Division of Water, Environment and Forestry Technology, Pretoria, South Africa. https://www.academia.edu/download/66385091/Empowering_communities_to_manage_natural20210420-1792-11yqufd.pdf

Sigalla, O. Z., Valimba, P., Selemani, J. R., Kashaigili, J. J., & Tumbo, M. (2023). Analysis of spatial and temporal trend of hydro-climatic parameters in the Kilombero River Catchment, Tanzania. Scientific Reports, 13(1), 7864.

Songorwa, A. N. (1999). Community-based wildlife management (CWM) in Tanzania: Are the communities interested? World Development, 27(12), 2061–2079.

Songorwa, A. N., Bührs, T., & Hughey, K. F. (2000). Community-based wildlife management in Africa: A critical assessment of the literature. Natural Resources Journal, 603–643.

Tang’are, J., & Mwanyoka, I. (2023). Assessing Factors Influencing Local Communities’ Compliance with Wildlife Conservation Regulations in Tanzania: A Case of Burunge Wildlife Management Area. Tanzania Journal of Forestry and Nature Conservation, 92(1), 214–229.

Tong, A., Sainsbury, P., & Craig, J. (2007). Consolidated criteria for reporting qualitative research (COREQ): A 32-item checklist for interviews and focus groups. International Journal for Quality in Health Care, 19(6), 349–357.

Treue, T., & Nathan, I. (2007). Community-based natural resource management. http://erepository.uonbi.ac.ke/handle/11295/42598

Tyrrell, P., Du Toit, J. T., & Macdonald, D. W. (2020). Conservation beyond protected areas: Using vertebrate species ranges and biodiversity importance scores to inform policy for an east African country in transition. Conservation Science and Practice, 2(1), e136. 10.1111/csp2.136

United Republic of Tanzania for Ministry of Natural Resources and Tourism. (1998). The Wildlife Policy of Tanzania. https://www.tnrf.org/files/E-URT_POLICIES_Wildlife_Policy_of_Tanzania_1998.pdf

United Republic of Tanzania for Ministry of Natural Resources and Tourism. (2012). The Wildlife Conservation (Wildlife Management Areas) regulations.

United Republic of Tanzania for Ministry of Natural Resources and Tourism. (2023). *National Wildlife Management Area Strategy*. https://www.honeyguide.org/wp-content/uploads/2023/06/THE-NATIONAL-WILDLIFE-MANAGEMENT-AREAS-STRATEGY-NWMAS-2023-2033.pdf

USAID. (2013). Tanzania wildlife management areas evaluation: Final evaluation report (pp. 1–132). Tetra Tech ARD and Maliasili Initiatives. https://pdf.usaid.gov/pdf_docs/pdacy083.pdf

USAID. (2016). Analysis of WMA Financial Viability and Options Study. https://pdf.usaid.gov/pdf_docs/PA00TZ99.pdf

Velund, H. E. (2009). Dry season distribution and density of Puku antelope (kobus vardoni) in the Kilombero valley floodplain FLOODPLAIN FORDELING OG TETTHET AV PUKUANTILOPE (KOBUS VARDONI) I TØRKEPERIODEN PÅ. https://static02.nmbu.no/mina/studier/moppgaver/2009-Velund.pdf

Walsh, K. (2023). Blood host preferences and competitive inter-species dynamics within an African malaria vector species complex inferred from signs of animal activity around aquatic larval habitats distributed across a gradient of fully domesticated to fully pristine ecosystems in southern Tanzania. https://cora.ucc.ie/server/api/core/bitstreams/c4075a66-6929-43e9-8807-eb4a3488bb55/content

Walsh, K. A., Kavishe, D. R., Duggan, L. M., Tarimo, L. J., Msoffe, R. V., Manase, E., Govella, N. J., Eichhorn, M. P., Kaindoa, E. W., & Butler, F. (2024). Blood host preferences and competitive inter-species dynamics within an African malaria vector species complex inferred from signs of animal activity around aquatic larval habitats. bioRxiv, 2024–08.

Wilfred, P. (2010). Towards Sustainable Wildlife Management Areas in Tanzania. Tropical Conservation Science, 3(1), 103–116. 10.1177/194008291000300102

Wright, V. C. (2017). Turbulent terrains: The contradictions and politics of decentralised conservation. Conservation and Society, 15(2), 157–167.

Xu, W., & Butt, B. (2024). Rethinking livestock encroachment at a protected area boundary. Proceedings of the National Academy of Sciences, 121(38), e2403655121. 10.1073/pnas.2403655121

