## Appendix 1 discussion guide tool for "Stakeholder perspectives on the effectiveness of the Ifakara-Lupiro-Mang’ula Wildlife Management Area in Southern Tanzania"

| 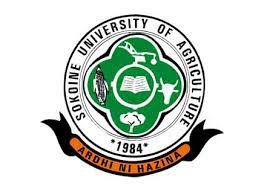 | **Comparing the perspectives of different stakeholders in the**  **Ifakara-Lupiro-Mang’ula**  **Wildlife Management Area** | 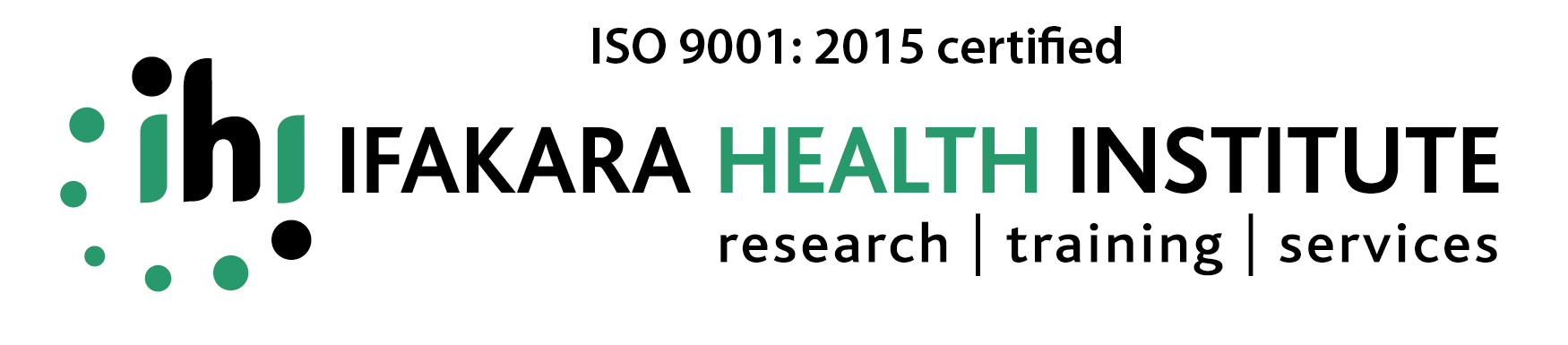 |
| --- | --- | --- |

**Discussion guide for FGDs and IDIs**

| **Interview date** |
| --- |
| **District** |
| **Ward** |
| **Village** |
| **Group/ Interviewee** |
| **Facilitator** |
| **Notes taker** |

**Respondent personal information**

| **ID no.** | **Age** | **Sex** | **Education** | **Years at current job** | **Years at current residence** |
| --- | --- | --- | --- | --- | --- |
| **1.** |  |  |  |  |  |
| **2.** |  |  |  |  |  |
| **3.** |  |  |  |  |  |
| **4.** |  |  |  |  |  |
| **5.** |  |  |  |  |  |

**SECTION ONE: GENERAL PERSPECTIVES ON THE ILUMA WMA**

1. Please explain briefly about the background information of ILUMA WMA.
2. What is your role regarding ILUMA Wildlife Management Area (ILUMA WMA)?
3. Please explain, to the best of your understanding, the reasons why the ILUMA WMA was established?
4. Do you think the establishment of ILUMA has had any beneficial or detrimental impacts? If yes or no, or a mixed picture, please describe and explain why.
5. How well do you think the livelihood support and community development functions of ILUMA WMA have worked? Please explain why.
6. How well do you think the environmental and wildlife conservation functions of ILUMA WMA have worked? Please explain why.
7. In which ways do people use land or other natural resources inside the conservation area of ILUMA WMA, with or without permission, and how does that affect wildlife, the environment, or other people in local stakeholder communities?
8. How well do you think the conservation functions of ILUMA complement its livelihood support and community development functions and *vice versa*? Or do they conflict with each other sometimes? Please explain why.
9. What is your view on the best way forward for the ILUMA WMA in the future? New ideas are especially welcome.

**SECTION 2. QUESTIONS FOR SOLICITING MORE IN-DEPTH PERSPECTIVES**

***Note to interviewer: These questions are optional and should be asked only if you feel they may be useful with respect to the respondent.***

**In relation to topic 4 about livelihoods and community development:**

4.1 How have livelihoods among local stakeholder communities been influenced by the WMA and why?
4.2 How have facilities, services and available local stakeholder communities been affected by the WMA and why?
4.3 How has the WMA influenced societal integrity, harmony, and representation among local stakeholder communities and why?
4.4 How much revenue do you think the ILUMA WMA has received from investors, researchers, tourists, hunting companies, resource use permits, fines and other sources since the new boundaries were agreed and established in 2014?
4.5 Please explain how much you know about how these ILUMA revenues are managed, who manages them and how well they are managed?
4.6 How effectively do you think these funds are used to benefit local communities?
4.7 Also, how effectively do you think these funds are used to ensure the protection of the environment and wildlife within the WMA conservation area?
4.8 Do you think that the benefits to local communities and investments in protecting the conservation area are adequate with respect to the revenues that ILUMA WMA generates?
4.9 What do you think could be done to ensure ILUMA WMA can generate more income, manage its income more effectively and spend it more effectively in the future?

**In relation to topic 5 about environmental and wildlife conservation functions**:

5.1 What is your view on the current conservation status of the environment and wildlife in ILUMA WMA compared to when the boundaries of the protected area that were agreed with the surrounding communities in 2014?
5.2 Also, if you are familiar with the ILUMA WMA since before establishment of the current boundaries, could you please share your views on how its longer-term history has influenced its status as a conservation area?
5.3 What attitudes do you think the local communities have towards the conservation functions of the ILUMA WMA?

Have they changed in any way since agreement and demarcation of the conservation area boundaries?

Please explain why you think any of these changes, for the better or for the worse, have occurred?

5.4 Looking ahead, what do you think are the challenges that ILUMA WMA faces with respect to conservation enforcement activities now and in the future?
5.5 Looking ahead, what is your opinion on improving the conservation functions of ILUMA WMA to make it more effective and sustainable?

**In relation to question 6 about how people use land or other natural resources inside the conservation area of ILUMA WMA, with or without permission:**

6.1 What do you think are the reasons for illegal use of land and other natural resources inside the conservation area of ILUMA WMA?
6.2 Please tell us how such illegal use of land and other natural resources inside ILUMA WMA are regulated, controlled, and prevented
6.3 Please describe how effective you think these regulations and activities to enforce them are?
6.4 How might these regulations and enforcement activities be improved upon, replaced, or supplemented to achieve more effective, sustainable, and acceptable conservation practice?
