## Appendix 2 participant consent form for "Stakeholder perspectives on the effectiveness of the Ifakara-Lupiro-Mang’ula Wildlife Management Area in Southern Tanzania"

| 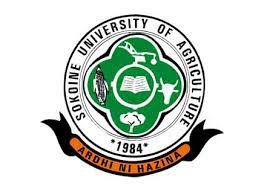 | **Comparing the perspectives of different stakeholders in the**  **Ifakara-Lupiro-Mang’ula**  **Wildlife Management Area** | 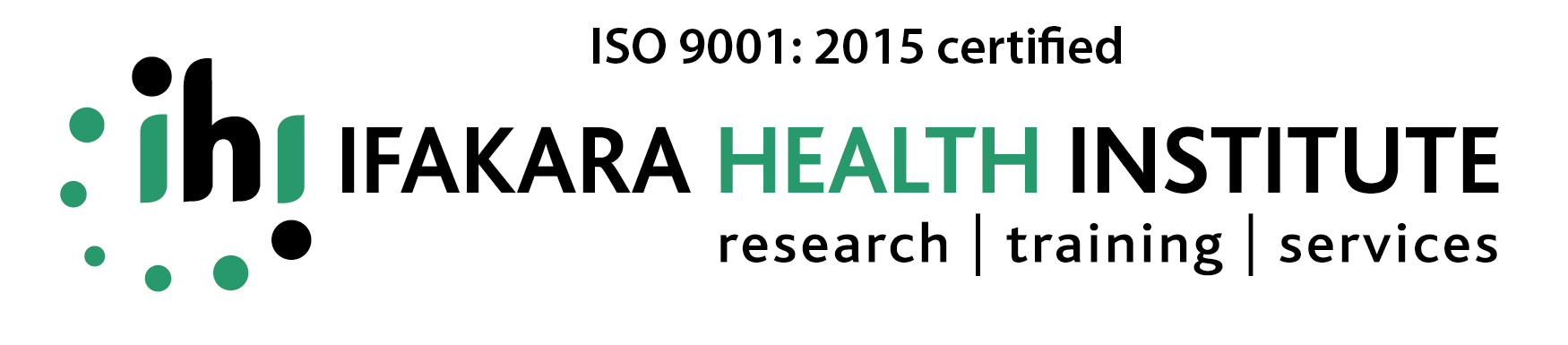 |
| --- | --- | --- |

**Participant consent form for WMA stakeholders**

In this study, we aim to compare the perspectives of different stakeholders on the conservation activities and contributions of ILUMA WMA to local livelihoods and community development.

We, therefore, invite you to participate in this study, so that you can share the perspectives you have as a stakeholder in the ILUMA WMA. We will need to record all the conversations we have with you so that later we can extract and describe all the detailed information you have kindly provided. Your participation in this study is entirely voluntary, and we would like to listen to your views on this topic. You may leave this study and end the conversation at any time, and you will not have to give any reason for doing so.

We don’t expect any major risks to you to arise from participating in this study, and we will keep all the information you provide in strict confidence. If we wish to quote anything that you said in subsequent presentations, publications, or other dissemination formats, we will not reveal your identity. If explaining your role is important background information for framing your perspective, and it might be possible to identify you individually based on that information, we will seek your permission in writing before doing so and will respect your wishes if you would prefer that quote not to be disseminated.

You will receive an allowance of Tsh 10,000/= for your participation in this study to compensate you for your time, meals, and inconvenience. Otherwise, no direct benefits are expected to arise from participating in this study. However, your participation in this study will be of great help to all the stakeholders in their effort to develop the ILUMA WMA.

All the information collected in this study will be kept strictly confidential, with paper hard copies stored in locked filing cabinets while soft copy electronic data will be stored on an encrypted password-protected computer and on an encrypted, password-protected server. Your personal information and the perspectives you share with us will only be used by the named researchers involved in this study and will not be disclosed to any other person. If there is any number of questions you may wish to ask, please feel free to ask them now or at any stage before, during, or after this recorded discussion.

The responsible scientist for this study is Ms. Lucia John Tarimo, a Research Assistant at the Department of Forest and Environmental Economics at the Sokoine University of Agriculture who is based at the Ifakara Branch of the Ifakara Health Institute, where it is overseen as a component of a larger study entitled “Population-stabilizing portfolio effects of fine-scale environmental variations in natural resource availability to malaria vector mosquitoes: characterization and implications for control strategies”. Correspondingly, the ethical aspects of this study involving human participants like yourself have been reviewed and approved by the Institutional Review Board of the Ifakara Health Institute.

If you have any questions related to ethics about this research please contact Ms.Lucia Tarimo at +255 763 006113 or Email address: or Prof. Felister Mombo at the Sokoine University of Agriculture at +255 785 252550 or Email address: or Mr. Deogratius Kavishe at the Ifakara Health Institute at +255 752 440654 or Email address:. In case you need further information, you may contact Dr. Mwifadhi Mrisho, the secretary of the Ifakara Health Institute ethical review board at +255 788 766676 or Email address:, or you may contact the secretary of National Health Research Ethics Review Committee (NatHREC) at +255 758 587885.

**Consent of the participant:**

I have read and understand all the information provided about this study. I voluntarily agree to participate in this study.

Name of the participant: ………………………………………………………………………….

Signature: …………………………………………………………………………………………

Date: ………………………………………………………………………………………………

**For an illiterate participant:**

A literate witness must sign (if possible, this person should be selected by the participant and should have no connection to the research team). Illiterate participants should include their thumbprints as well.

I have witnessed the accurate reading of the consent form to the potential participant, and the individual has had the opportunity to ask questions. I confirm that the individual has given consent freely.

Name of the witness: ………………………...…………………………………………………...

Signature of the witness: …………………………………………………………………………

Date: ……………………………………………………………………………………………...

A thumbprint of the participant

**Declaration of the applicant for consent to the participant**

I confirm that I have carefully read and explained all the above information to the above-named signatory participant and have done my best to ensure the participant has fully understood the risks, inconveniences, protections, and benefits associated with participation in this study**.**

I confirm that the participant was provided with adequate opportunity to ask questions about this study and that any questions that were asked were answered accurately and to the best of my ability. I confirm that the participant was not forced to sign the consent form but did so with his or her fully informed consent.

A copy of the proof of consent to participate is provided to the participant.

Name of the person who requested consent to participate: ………………………….…………………….

Signature of the person who requested the consent to participate: ………………………………………..

Date: ……………………………………………………………………………………………………….
